# Molecular mechanism by which SARS-CoV-2 Orf9b suppresses the Tom70-Hsp90 interaction to evade innate immunity

**DOI:** 10.1101/2025.11.18.689095

**Authors:** Noah Sherer, Abhishek Bastiray, Xiao-Ru Chen, Trivikram Molugu, Gaya P. Yadav, Tatyana I. Igumenova, Jae-Hyun Cho

## Abstract

The Tom70-Hsp90 interaction is critical for activating MAVS-mediated interferon (IFN) production. Upon RNA virus infection, cytosolic Hsp90 recruits key innate immune signaling proteins to MAVS on mitochondria through its interaction with Tom70. To evade this innate immune response, the SARS-CoV-2 protein Orf9b binds to Tom70, thereby disrupting the Tom70-Hsp90 interaction and suppressing IFN production. Despite its importance, the molecular mechanism underlying Orf9b-mediated inhibition of IFN signaling remains unclear. Here, using an integrative approach combining cryo-electron microscopy, ^19^F NMR spectroscopy, and isothermal titration calorimetry (ITC), we show that Orf9b inhibits Hsp90 binding to Tom70 through a bipartite mechanism. The helix and intrinsically disordered tail of Orf9b sterically block the access of two distinct structural units of Hsp90 to Tom70. We also find that Orf9b-mediated allosteric conformational changes in Tom70 do not contribute to the inhibition of the Hsp90 binding. Comprehensive structural, thermodynamic, and kinetic analyses further reveal that Orf9b primarily slows the association kinetics between Hsp90 and Tom70. Collectively, our results provide a high-resolution mechanistic framework for understanding Orf9b-mediated suppression of the host innate immune response.

## Introduction

Mitochondria play a central role in interferon expression which is the hallmark of innate immune responses during viral infection^1,2^. Upon detection of viral dsRNA by RIG-I-like Receptors (RLRs), a number of proteins are recruited to the mitochondrial surface, forming the mitochondrial antiviral signaling-associated (MAVS) signalosome leading to interferon expression^3–6^. Translocase of outer mitochondria membrane 70 (Tom70) mediates the interaction of MAVS with downstream effectors (**Fig 1A**). For example, Tom70 acts as a receptor for heat shock protein 90 (Hsp90) that recruits TANK-binding kinase 1 (TBK1) and interferon regulatory factor 3 (IRF3) to MAVS^7,8^. Therefore, the Tom70-Hsp90 interaction is important for establishing innate immune responses upon viral infection^9^. Moreover, the Tom70-Hsp90 interaction mediates transferring cytosolic preproteins into mitochondria, which is important for mitochondrial biogenesis^10–13^.

**Figure 1:**
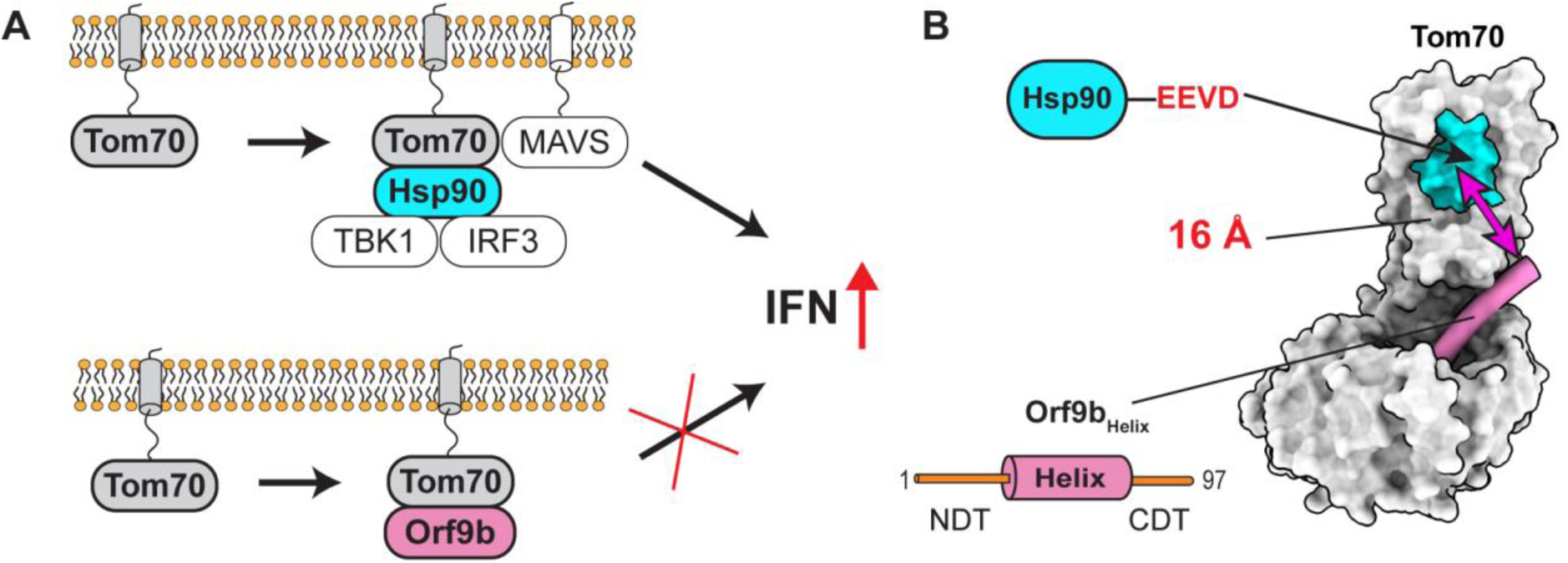
SARS-CoV-2 Orf9b inhibits Tom70 mediated IFN-I production. **(A)** The Tom70-mediated interferon response without (top) and with (bottom) SARS-CoV-2 Orf9b. Orf9b interferes with the Tom70-Hsp90 interaction, resulting in suppression of IFN production. **(B)** Surface representation of the Tom70-Orf9b complex (PDB: 7DHG). The putative Hsp90 binding site (cyan) is located more than 16Å away from the visible part of the bound Orf9b (pink). The NDT and CDT of Orf9b represent the N- and C-terminal disordered tails (orange), respectively, and they were not visible in the structure.

Mitochondrial dysfunction has been identified in acute and long COVID-19 patients^14,15^. Open reading frame 9b (Orf9b) is considered one of major virulence factors of SARS-CoV-2^16,17^. Orf9b directly binds to Tom70, inhibiting Hsp90 binding to Tom70 and subsequently IFN expression (**Fig 1A**)^17–19^. Hence, inhibition of the Tom70-Hsp90 interaction by Orf9b is a critical step to suppress the host innate immune response during infection of SARS-CoV-2^17,20,21^, including variants of concerns^16^. Despite its importance, the molecular basis of how Orf9b inhibits the Tom70-Hsp90 interaction remained unclear.

Previous cryo-EM and crystal structures of the Tom70-Orf9b complex showed only a portion of Orf9b bound to hydrophobic pocket of Tom70, while the rest was invisible due to flexibility (**Fig 1B**)^17,18^. The visible part of Orf9b forms an α-helix that is distant (> 16 Å) from the putative Hsp90-binding site on Tom70 (**Fig 1B**). This observation led to the proposal of an allosteric inhibition model. In that model, Orf9b binding to Tom70 induces allosteric conformational changes at the Hsp90 binding site that reduce the affinity of Tom70-Hsp90 interactions^17,18^. However, the allosteric model remained untested due to the lack of structural information on human Tom70 (hTom70) in its free and Hsp90-bound states, which is essential for assessing whether Orf9b binding leads to allosteric conformational changes at the Hsp90-binding site.

The structural basis of the Tom70-Hsp90 interaction has so far been studied only for yeast proteins^22,23^. However, human and yeast Tom70 and Hsp90 share only 20% (68%) and 60% (87%) amino acid identity (similarity), respectively (**Figs S1 and S2**). In addition, some structural and functional differences between mammalian and yeast Tom70 (yTom70) have been reported^13,24,25^. Notably, yeast does not have an innate immune system, such as for IFN expression^26,27^. Consequently, the mechanistic basis and extent of Orf9b-mediated inhibition of the human Tom70-Hsp90 interaction remains unknown.

To determine how Orf9b inhibits the Tom70-Hsp90 interaction, we employed an integrative approach that combines cryo-EM, ^19^F NMR spectroscopy, ITC, and BLI for structural, thermodynamic, and kinetic studies. Briefly, we determined the structures of human Tom70 in its free and Hsp90-bound states. We also examined the thermodynamic and kinetic effects of Orf9b on the Tom70–Hsp90 interaction. Surprisingly, we did not observe Orf9b-mediated allosteric inhibition of the Tom70–Hsp90 interaction, despite minor allosteric conformational changes. Instead, our comprehensive analysis revealed that Orf9b inhibits Hsp90 binding through a bipartite steric exclusion mechanism involving the helix and the C-terminal disordered tail of Orf9b. Furthermore, we found that this inhibition arises from reduced association kinetics of Hsp90 with Tom70 due to an entropic bristle effect by the disordered tail of Orf9b. Collectively, these findings provide a new mechanistic basis for the Orf9b-mediated immune evasion strategy of SARS-CoV-2.

## Results

### Effects of Orf9b on the Tom70-Hsp90 interaction

Although Orf9b is known to inhibit the Tom70-Hsp90 interaction, its inhibitory effect has not been quantitatively assessed. The lack of this essential information has hampered understanding the mechanism of Orf9b-mediated inhibition of the Tom70-Hsp90 interaction. Thus, we measured the changes in the Tom70-Hsp90 affinity in the absence and presence of Orf9b using a series of ITC experiments.

The binding affinity (K_D_) of full-length Hsp90 to free Tom70 was 3 ± 1 µM, which falls within the affinity range of Hsp90 to its other clients^28–30^ (**Figs 2A and S3A**). To measure Orf9b’s inhibitory effect on the Hsp90-Tom70 interaction, we titrated the Tom70:Orf9b complex into Hsp90. The binding affinity of the Hsp90 to the Tom70:Orf9b complex was 22 ± 1 µM, representing a 7.3-fold reduction compared to binding to free Tom70 (**Figs 2B and S3B**). We further confirmed that the Tom70:Orf9b complex has an expected molecular weight (69 kDa) using SEC-MALS (**Fig S4A)**.

**Figure 2:**
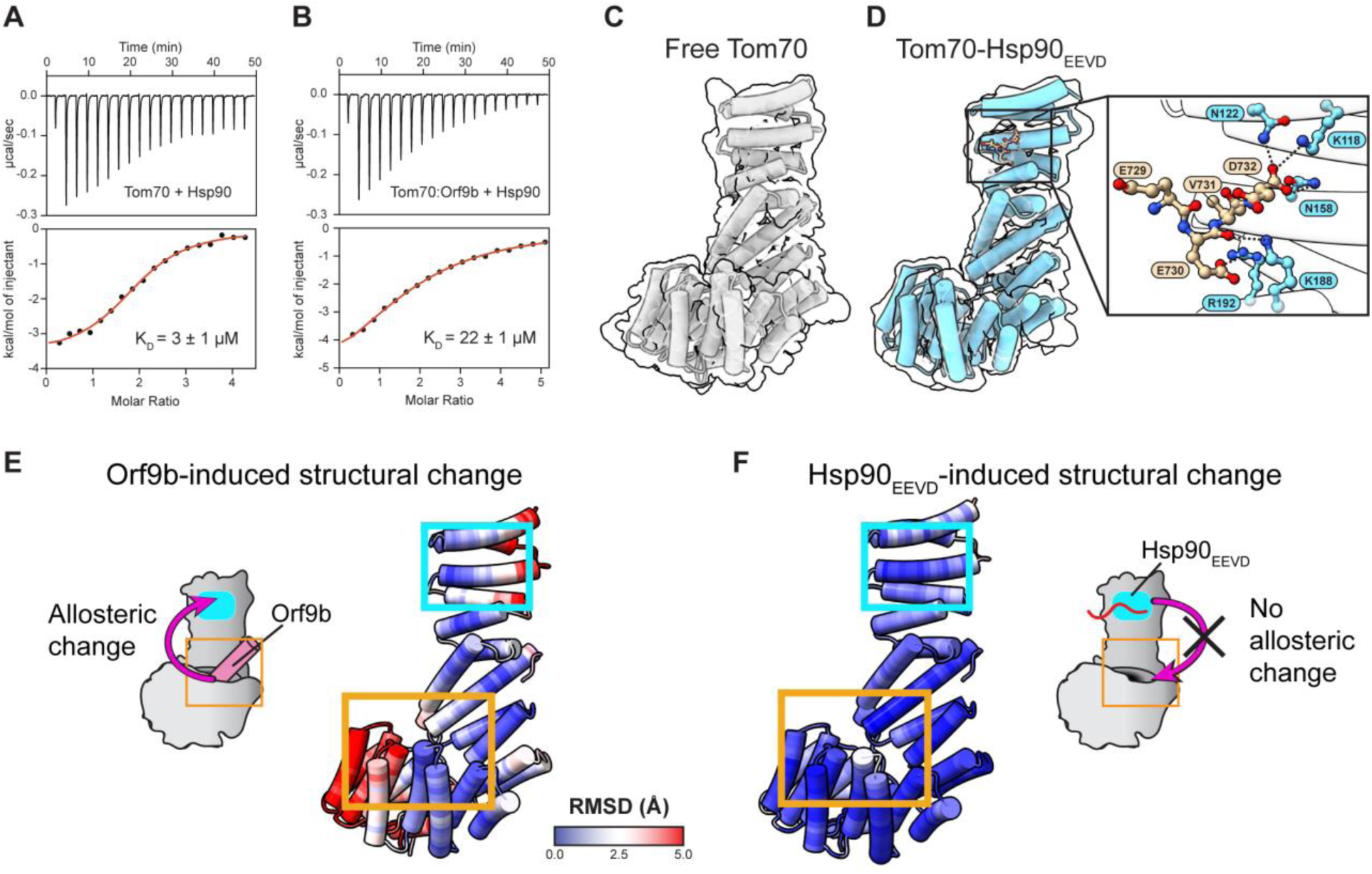
Cryo-EM structures of Tom70 and Tom70-Hsp90_EEVD_ reveal minimal structural allostery. (A-B) ITC thermograms and isotherms for the interaction between Tom70 and full-length Hsp90 in the **(A)** absence and the **(B)** presence of full-length Orf9b. The red line indicates the fit curve for a 1:1 binding model. Numbers after the ± symbol indicate the standard deviation of 3 measurements. **(C)** Cryo-EM map and model of free Tom70. **(D)** Cryo-EM map and model of Tom70 complexed with Hsp90_EEVD_ peptide. The inset shows the interactions between Tom70 and Hsp90_EEVD_. **(E-F)** RMSD analysis for free Tom70 vs **(E)** Orf9b-bound (PDB: 7DHG) and **(F)** Hsp90_EEVD_-bound. RMSD values are mapped onto free Tom70.

Hsp90 binds to Tom70 using its C-terminal EEVD motif^31–33^. Hence, to investigate how Orf9b inhibits the Tom70–Hsp90 interaction, we determined cryo-EM structures of free Tom70 (**Fig 2C)** and the Hsp90 EEVD peptide (Hsp90_EEVD_)-bound hTom70 (**Fig 2D) (Figs S5 and S6, and Table S1**). Together with the previous hTom70-Orf9b structure, these structures enabled detailed analysis of Orf9b- and Hsp90_EEVD_-mediated conformational changes in Tom70.

In the Tom70:Hsp90_EEVD_ complex, electron density for the Hsp90_EEVD_ was observed only for the C-terminal four residues out of the total 10, indicating a lack of interactions with Tom70 other than the four residues. These visible residues formed specific salt-bridge interactions with hTom70 (**Fig 2D and S7A**). These interactions were further confirmed by site-directed mutagenesis (**Fig S7B-C**).

Briefly, the C-terminal α-carboxyl group of the Hsp90_EEVD_ peptide forms a salt bridge and hydrogen bond with K118 and N158 of hTom70 respectively (**Fig 2D**). A synthetic peptide with an amidated C-terminus completely abolished binding to Tom70, and K118A and N158A mutations in hTom70 also eliminated binding (**Fig S7B and S7C**). Another salt bridge and hydrogen bond were identified between E728 in the Hsp90_EEVD_ motif and R192 and K188 in Tom70 respectively. Both the K188A and the R192A mutation likewise abrogated binding (**Fig S7B and S7D)**.

Next, we assessed potential conformational changes in hTom70 due to Orf9b binding. Structural differences, measured as root mean square deviations (RMSD) between free and Orf9b-bound Tom70, revealed substantial conformational changes in the hydrophobic pocket in the C-terminal domain of Tom70 due to direct interactions with bound Orf9b (**Fig 2E**). Specifically, the α-helices forming the hydrophobic pocket shifted outward to accommodate Orf9b. In contrast, the N-terminal domain of Tom70, which harbors the Hsp90-binding site, showed relatively minor changes upon Orf9b binding. Moreover, all residues involved in Hsp90 binding remained well exposed to solvent in both free and Orf9b-bound hTom70 (**Figs S8A and S8B)**. Consistently, the pairwise distances between Hsp90_EEVD_ and hTom70 showed no significant changes in the Tom70-Orf9b complex compared to those in free Tom70 (**Fig S8C**).

We further examined whether Hsp90 binding to Tom70 causes reciprocal conformational changes at the Orf9b binding site. Yet, structures of free Tom70 and Tom70:Hsp90_EEVD_ complex showed only minimal changes in the Orf9b-binding pocket upon Hsp90 binding (**Fig 2F**). Collectively, these results suggest that Orf9b-mediated inhibition of the Tom70–Hsp90 interaction is unlikely to arise from allosteric conformational changes.

### Thermodynamic cycle analysis reveals the absence of allosteric inhibition

To ascertain that Orf9b is not an allosteric inhibitor of the hTom70-Hsp90 interaction, we designed a minimalist model consisting of hTom70, the Hsp90_EEVD_ peptide, and the Orf9b peptide containing only the helical region (Orf9b_Helix_) observed in the Tom70–Orf9b complex (**Fig 3A**). Importantly, due to their small size, these minimal peptides do not directly contact each other, even when both are bound to Tom70 simultaneously. This feature makes the system ideal for probing allosteric effects without direct contact between the binding partners.

**Figure 3:**
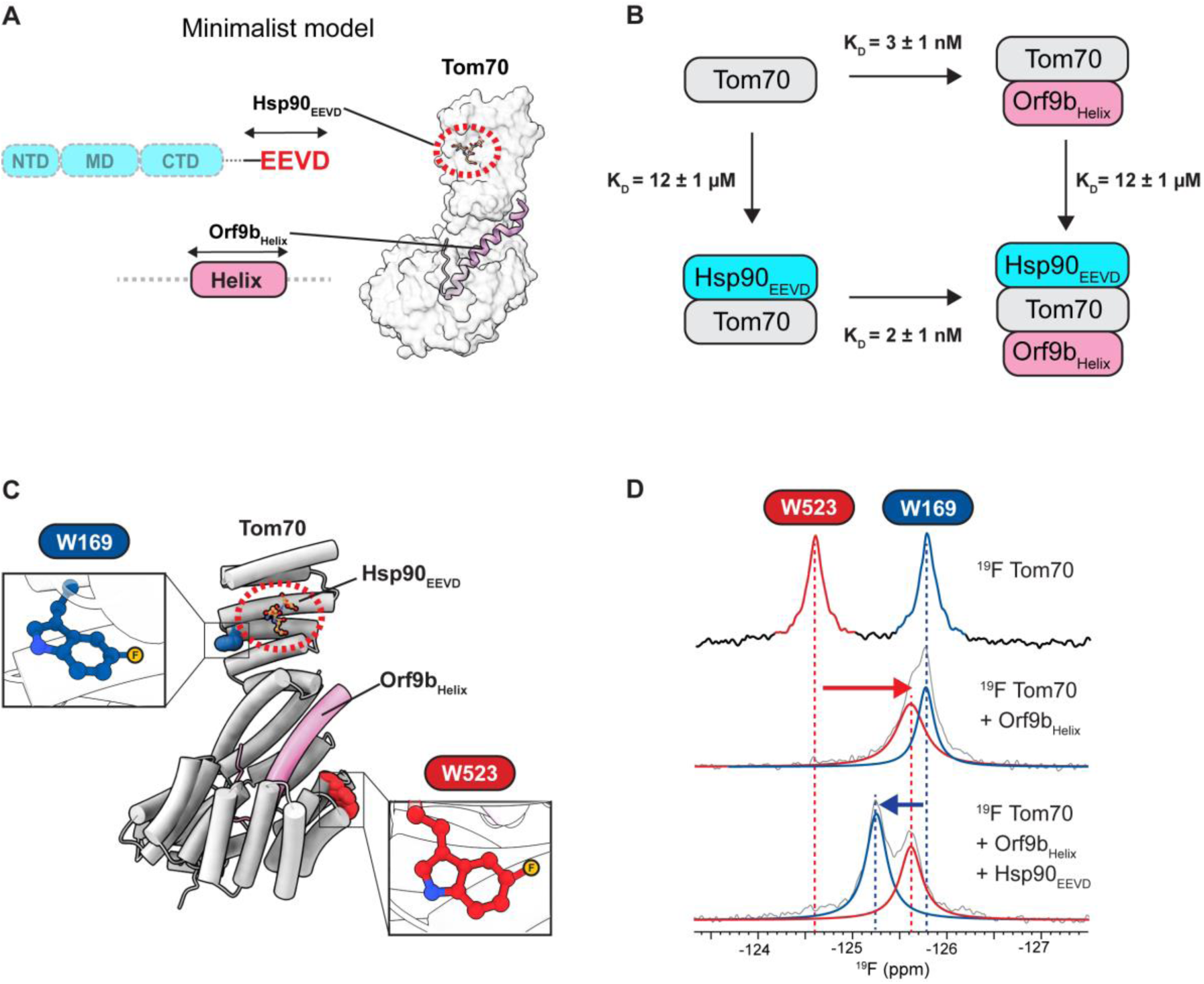
Orf9bHelix does not allosterically inhibit Hsp90_EEVD_ binding to Tom70. **(A)** Schematic of minimalistic system to test for allostery. Orf9b_Helix_ contains only the residues visualized in the crystal structure bound to Tom70 (PDB: 7DHG). Hsp90_EEVD_ is a 10-residue peptide consisting of the C-terminal residues of Hsp90. **(B)** Thermodynamic cycle analysis for Orf9b_Helix_ and Hsp90_EEVD_ binding to Tom70. Numbers after the ± symbol indicate the standard deviation of 3 measurements. **(C)** Structure of Tom70 artificially superimposed with Hsp90_EEVD_ and Orf9b_Helix_. Two 5F-Trp residues are shown in a sphere model. Insets show close-up views of the two Trp residues, with the 5F positions highlighted as a yellow ball. **(D)** ^19^F NMR spectra of Tom70 in the free (top), Orf9b_Helix_-bound (middle), and Orf9b_Helix_- and Hsp90_EEVD_-co-bound (bottom) states. Gray lines represent the observed ^19^F spectra. Blue and red lines represent the deconvoluted signals of W169 and W523, respectively.

We next conducted a comprehensive thermodynamic cycle analysis^34^ to test the allosteric inhibition model. Interestingly, we found that the ITC-derived binding affinities of Hsp90_EEVD_ for free Tom70 and for the Tom70:Orf9b_Helix_ complex were virtually identical, indicating that the Orf9b_Helix_ has no measurable effect on the EEVD motif-Tom70 interaction (**Figs 3B, S9A, and S9B**). Likewise, we also observed that Hsp90_EEVD_ does not inhibit Orf9b_Helix_ binding to Tom70 (**Figs 3B, S9C, and S9D)**.

These results refute the allosteric inhibition model. Although surprising, it is consistent with our structural analysis showing that Hsp90-binding site remains accessible in the Tom70-Orf9b structure (**Figs 2E, 2F, S8B, and S8C**). It is also noteworthy that the Hsp90_EEVD_ motif resides in the long-disordered C-terminal tail, which could readily undergo conformational adjustments to bind Tom70, as observed in the yTom71-Hsp70/90_EEVD_ complexes^23,35^ (**Fig S10A**).

To further assess potential Orf9b-mediated conformational changes in Tom70 in solution, we employed ^19^F NMR spectroscopy, which is exquisitely sensitive to local structural changes and greatly simplifies the spectral complexity of large proteins^36–39^. Notably, Tom70 contains only two tryptophan residues, W169 and W523, located next to the EEVD-binding site and the Orf9b-binding pocket, respectively (**Fig 3C**). The indole rings of both W169 and W523 are solvent exposed, so incorporation of fluorinated tryptophan is unlikely to perturb the structure. These properties make ^19^F NMR an ideal method to probe possible allosteric changes in Tom70 upon Orf9b or Hsp90 binding.

We incorporated 5F-tryptophan (^19^F at position 5 of the indole ring) into Tom70, which yielded two distinct ^19^F NMR signals corresponding to 5F-W169 and 5F-W523 (**Fig 3D**). Upon addition of Orf9b_Helix_, only the 5F-W523 peak shifted, whereas the 5F-W169 peak remained unchanged. Conversely, when Hsp90_EEVD_ was added to the Tom70:Orf9b_Helix_ complex, only the 5F-W169 peak shifted, with no change in the position of the 5F-W523 peak at the Orf9b-binding site (**Fig 3D**). These results demonstrate that Orf9b_Helix_ and Hsp90_EEVD_ peptides can co-bind to hTom70, forming a ternary complex, which is consistent with our ITC results.

We further examined the effect of full-length Orf9b and Hsp90 onto the ^19^F NMR spectrum of hTom70. Remarkably, binding of full-length Orf9b and Hsp90 also led to changes only in the peak positions of 5F-W523 and 5F-W169, respectively, without affecting the other site (**Fig S11**). These results are the same as those observed with the minimalist model, supporting the conclusion that no significant conformational changes occur that could lead to allosteric inhibition. However, it should be noted that these results do not indicate the absence of Orf9b-mediated inhibition of the Tom70-Hsp90 interaction. Our ITC data clearly shows that full-length Orf9b inhibits Hsp90 binding (**Figs 2A and 2B**). Rather, these results point to a non-allosteric mechanism of inhibition.

### Bipartite inhibition mechanism mediated by the Orf9b disordered tail and helix

Given these findings, we set out to identify an alternative inhibition mechanism. In this light, we noted that the inhibitory effect of full-length Orf9b exceeds that of the Orf9b_Helix_ by 2.4-fold (**Figs 2B, 4A, 4B, S3B, and S3C**). Because the Orf9b_Helix_ lacks the N- and C-terminal disordered tails, this result suggested that the Orf9b-mediated inhibition arises from two distinct but cooperative parts, i.e., the helix and disordered tails of Orf9b.

**Figure 4:**
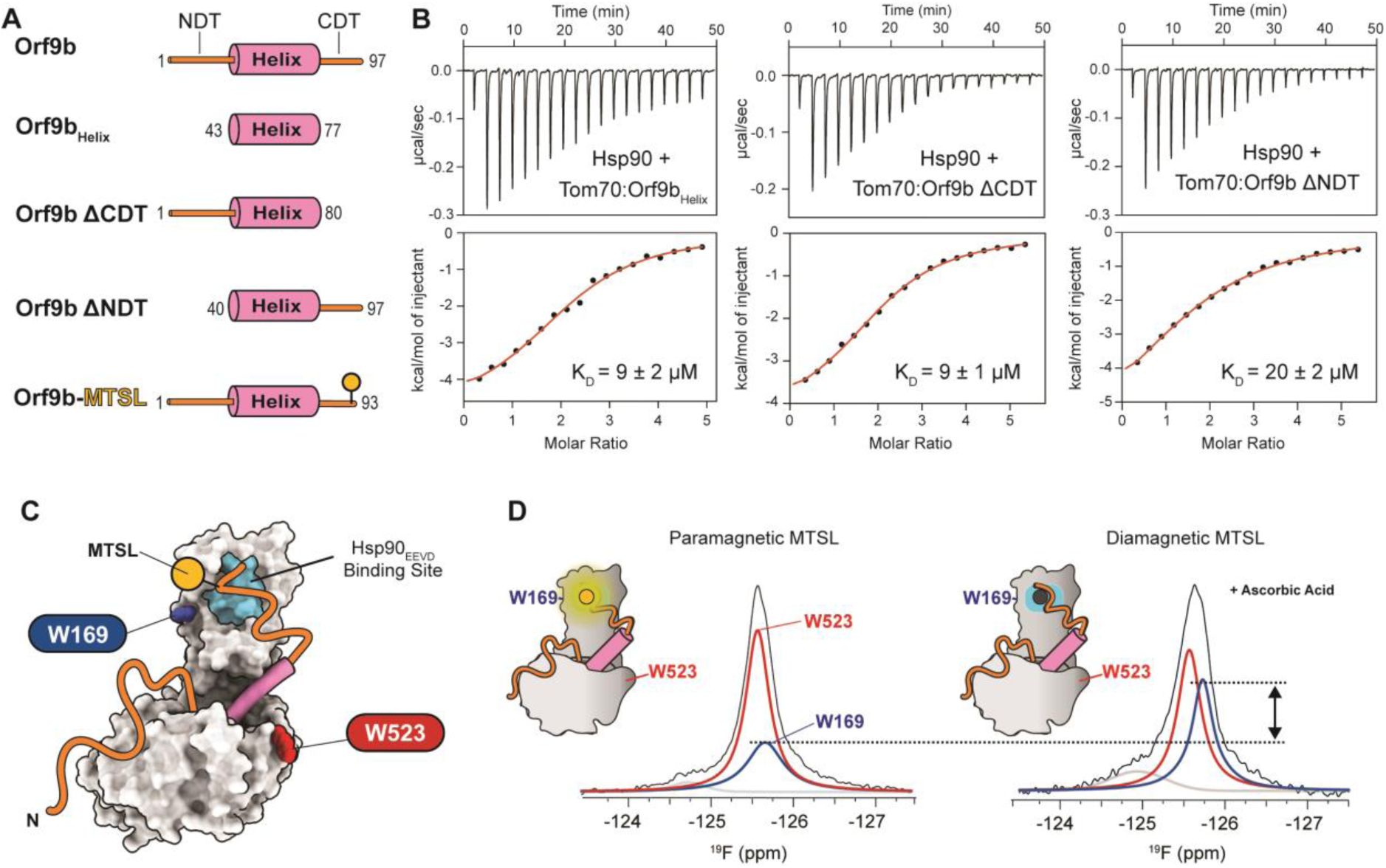
The Orf9b C-terminal disordered tail inhibits Hsp90 binding to Tom70. **(A)** Schematic of all Orf9b variants. **(B)** ITC thermograms and isotherms for the interaction between Tom70 and Hsp90 in the presence of Orf9b_helix_ (left), Orf9b ΔCDT (middle), and Orf9b ΔNDT (right). Numbers after the ± represent the standard deviation of at least two measurements. **(C)** Schematic of the PRE experiment. The MTSL spin label (yellow ball) is attached at the CDT of Orf9b. **(D)** ^19^F spectra of Tom70 complexed with Orf9b labeled with paramagnetic (left) and diamagnetic (right) MTSL. Black lines represent the observed ^19^F spectra. Blue and red lines represent the deconvoluted ^19^F peaks corresponding to W169 and W523, respectively.

First, to understand how the disordered tails of Orf9b inhibit the Tom70-Hsp90 interaction, we examined which of the N- and C-terminal disordered tails (NDT and CDT respectively) blocks the binding of the Hsp90_EEVD_ motif to Tom70. We generated two truncated Orf9b variants: Orf9b ΔNDT and Orf9b ΔCDT, in which the NDT and CDT were deleted, respectively (**Fig 4A**). Binding affinities of Hsp90 to the Tom70–Orf9b ΔNDT and Tom70–Orf9b ΔCDT complexes were then measured by ITC (**Fig 4B, S3D, and S3E**). We found that Orf9b ΔCDT exhibited the same inhibitory effect as Orf9b_Helix_, whereas Orf9b ΔNDT showed a similar inhibitory effect as full-length Orf9b. These results demonstrate that the Orf9b CDT is primarily responsible for inhibiting Hsp90 binding to Tom70.

To confirm that the Orf9b CDT is spatially close to the EEVD binding site in the Tom70-Orf9b complex, we employed ^19^F NMR paramagnetic relaxation enhancement (PRE)^40–43^. A V92C mutation was introduced into Orf9b for labeling with MTSL, and the MTSL-labeled Orf9b was mixed with ^19^F-labeled Tom70 (**Fig 4C**). We observed a significant reduction in the peak intensity of ^19^F-W169 (**Fig 4D**), which is located near the Hsp90_EEVD_ binding site. As expected, the peak intensity increased upon reduction of MTSL with ascorbic acid. The ^19^F peak intensity ratio of W169 to W523 clearly indicated that the Orf9b CDT is spatially close to W169, although quantitative estimation of the PRE effect was not feasible due to minor but detectable dimerization of Tom70 in the absence of reducing agent (i.e., TCEP).

Next, we examined how the Orf9b helix contributes to the inhibition of Hsp90 binding. Although the structure of the full-length Hsp90-Tom70 complex is not yet available, we noted that the C-terminal domain (CTD) of Hsp90 is thought to form a secondary association near the hydrophobic pocket of Tom70^32,44^, where the Orf9b helix binds (**Fig 5A).** Hsp90 is a homodimer with a molecular weight of ∼180 kDa. Considering its large size, it is plausible that Hsp90 binding to this secondary site is sterically hindered by the helix of bound Orf9b. Indeed, we observed that full-length Hsp90 binds to free Tom70 with 4-fold higher affinity than the Hsp90_EEVD_ peptide (**Figs 2A, 3B, S3A, and S9A**). This result supports that full-length Hsp90 interacts with Tom70 through an additional non-EEVD interface that is disrupted by the bound Orf9b helix (**Fig 5B**).

**Figure 5:**
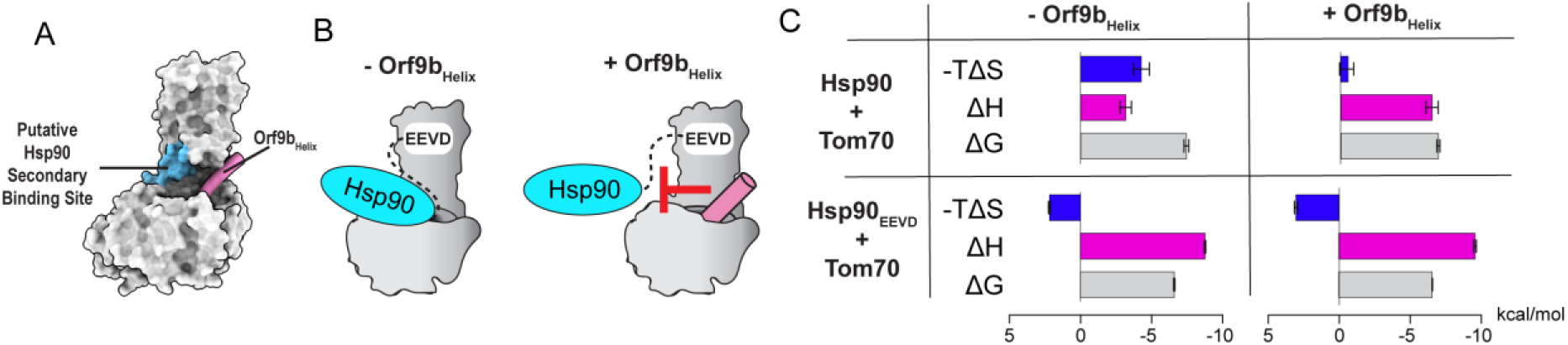
The Orf9b helix sterically prevents a secondary Hsp90 binding site. **(A)** Schematic showing the putative Hsp90 ancillary binding site (cyan) and Orf9b_Helix_ (pink) displayed on a surface representation of the Tom70-Orf9b structure (PDB: 7DHG). **(B)** Cartoon representation of how the Orf9b_Helix_ may sterically block the additional Hsp90 binding site. **(C)** ITC-derived thermodynamic signatures for full-length Hsp90 and Hsp90_EEVD_ binding to Tom70 in the absence and presence of Orf9b_Helix_. Error bars represent the standard deviation of at least three measurements.

Perturbation of the secondary Hsp90-Tom70 interaction by the Orf9b helix is further supported by changes in the thermodynamic signature of binding. Specifically, we observed that a favorable entropy change was the major driving force for full-length Hsp90 binding to free Tom70 (**Figs 5C and S3A**), suggesting that a large hydrophobic surface becomes buried upon complex formation^45–47^. In contrast, full-length Hsp90 binding to the Tom70-Orf9b_Helix_ complex was driven entirely by a favorable enthalpy change, indicating the loss of this hydrophobic binding interface^48^ (**Figs 5C and S3C)**. Unlike full-length Hsp90, however, the Hsp90_EEVD_ peptide binds to Tom70 with virtually identical thermodynamic signature in the absence and presence of Orf9b_Helix_ (**Figs 5C, S9A, and S9B**), excluding the possibility that Orf9b_Helix_ binding induces an undetected conformational change in the Hsp90_EEVD_-binding site. Taken together, these results strongly support the steric blocking model of Orf9b in which the CDT and helix prevent the access of Hsp90_EEVD_ motif and Hsp90_CTD_ to their respective Tom70 binding sites.

### Orf9b inhibits the association kinetics of Hsp90 with Tom70

If a protein–protein interaction is sterically blocked, the association rate constant (k_on_) is most likely to be affected^49^. We therefore examined whether the association rate of Hsp90 with Tom70 is inhibited by bound Orf9b, using BLI to measure binding kinetics. However, the association rate of Hsp90 to free Tom70 was too rapid to quantify accurately, although it was evident that the association rate was significantly reduced in the presence of Orf9b (**Figs 6A, S12A, and S12B**).

**Figure 6:**
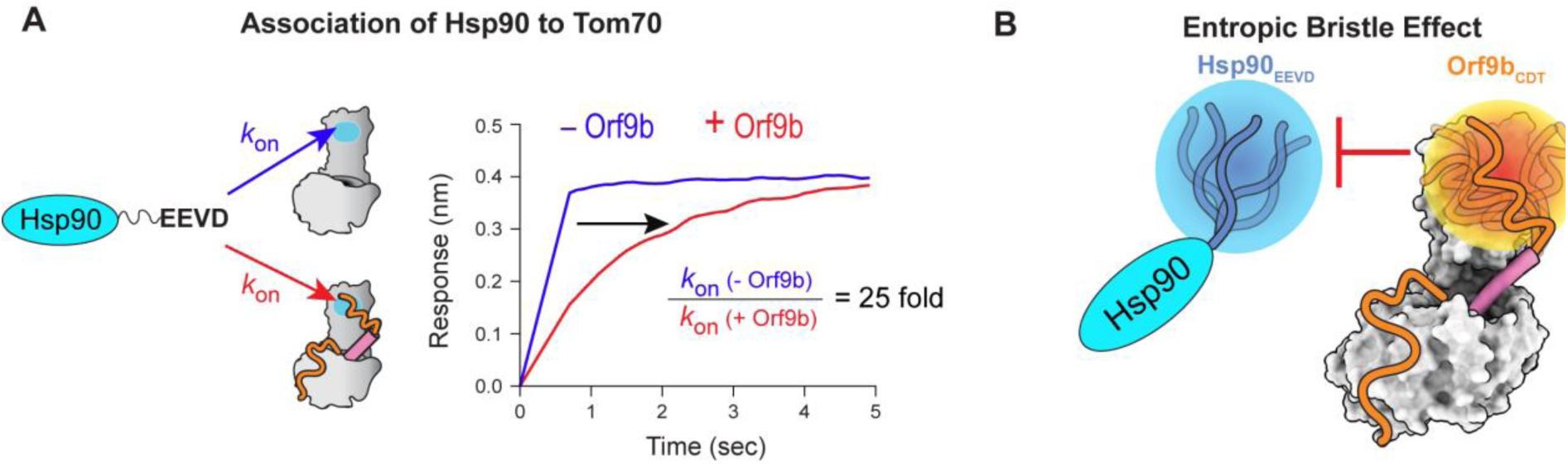
Orf9b acts as an entropic bristle to inhibit Hsp90 binding to Tom70. **(A**) (Left) Schematic illustrating the Hsp90_EEVD_-mediated association to Tom70 in the absence (blue arrow) and presence (red arrow) of Orf9b. (Right) Representative BLI sensorgrams showing the association of Hsp90 to Tom70 in the absence (blue) and presence (red) of Orf9b. **(B)** Schematic of the entropic bristle inhibition model. The conformational dynamics of the C-terminal disordered tail of Orf9b (Orf9b_CDT_) sterically hinders the Hsp90_EEVD_ motif from binding to Tom70. The blue and orange spheres represent hypothetical excluded volumes generated by the flexible C-terminal tails of Hsp90 and Orf9b, respectively.

By contrast, the dissociation rate constant (k_off_) could be measured reliably for the Tom70–Hsp90 complex in both the absence and presence of Orf9b (**Figs S12A and S12B**). We therefore indirectly estimated k_on_ values by combining k_off_ values obtained from BLI with K_D_ values determined by ITC, i.e., 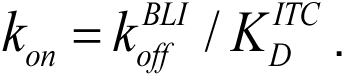 The only assumption was that relative differences in K_D_ values of Tom70-Hsp90 with and without Orf9b (ΔK_D_) are consistent across the two methods, i.e., ΔK_D_^ITC^ ≍ ΔK_D_^BLI^. This analysis revealed that the k_on_ value for Hsp90 binding to Tom70 (k_on_ = 0.5 ± 0.2 µM^-1^s^-1^) was reduced by approximately 25-fold in the presence of bound full-length Orf9b (k_on_ = 0.02 ± 0.01 µM^-1^s^-1^) (**Fig S12C)**. These results are consistent with the NMR and ITC results that the Orf9b CDT is primarily responsible for steric inhibition of the Hsp90_EEVD_ motif binding to Tom70 (**Fig 6B**).

## Discussion

In this study, we provide new mechanistic insights into Orf9b-mediated inhibition of the Tom70–Hsp90 interaction. We found that the structurally invisible C-terminal disordered tail of Orf9b is primarily responsible for inhibiting the association of the Hsp90 EEVD motif with Tom70. Structural disorder increases the radius of gyration of a protein, enabling it to effectively block access to the EEVD binding site that is also located in the disordered tail of Hsp90. This role of the disordered region of Orf9b can be described as an entropic bristle^50,51^, where intrinsically disordered regions prevent access of other proteins through their high flexibility (**Fig 6B**). The entropic bristle raises the free-energy barrier for association, however, once bound, k_off_ values often change little because the main penalty was entering the excluded volume of bristle layer, not staying in the complex^49,52,53^. Indeed, we observed the significantly reduced k_on_ of Hsp90 to Tom70 by Orf9b (25-fold change), with a minimal change in k_off_ values (3-fold change) **(Fig S12)**. Furthermore, the Orf9b_Helix_ sterically occludes a secondary Hsp90 interaction near the hydrophobic pocket of Tom70. Consequently, the Orf9b helix and the disordered tail together constitute the bipartite inhibition mechanism **(Fig S13)**.

The Orf9b_Helix_ has previously been proposed to induce allosteric conformational changes in the Hsp90-binding site on Tom70^17,18,44^. Although we also observed allosteric conformational changes in hTom70, the key Hsp90-binding residues remained accessible by the Hsp90_EEVD_ motif **(Figs S8B and S8C)**. Moreover, we found that the allosteric conformational change is non-reciprocal between the Orf9b_Helix_-binding and Hsp90_EEVD_-binding sites in hTom70 **(Figs 2E and 2F).** Most importantly, protein allostery is a thermodynamic phenomenon, i.e., binding at an allosteric site modulates binding affinity at another site, through diverse mechanisms^54–57^. Thus, our thermodynamic cycle analysis provides unambiguous evidence for the absence of allosteric inhibition by Orf9b **(Fig 3B)**. These results underscore that allosteric conformational change does not necessarily equate to allosteric inhibition^58^. We speculate that the allosteric conformational change in Tom70 might play a role in the transfer of mitochondrial preproteins^44^.

It is noteworthy that our ITC results differ from the values previously reported values by Gao et al^18^. Specifically, they reported a 13-fold reduction in the binding affinity of Hsp90_EEVD_ to Tom70 in the presence of the Orf9b_Helix_ peptide, which we did not observe (**Fig 3B**). Instead, we observed a notable reduction of the binding affinity only when full-length Orf9b and Hsp90 were employed in the ITC measurements. Our results are further supported by a comprehensive thermodynamic cycle analysis (**Fig 3B**), structural analysis using cryo-EM (**Figs 2D and 2E)**, and ^19^F NMR data **(Fig 3D)**. The source of this difference remains unclear. We have appended all our individual ITC results for comparison (**Figs S3 and S9)**.

The allosteric inhibition model was originally based on structural comparisons between human Tom70 (hTom70) and yeast Tom70 (yTom70)^22,23,35^, because structures of hTom70 in the free and Hsp90-bound states were not previously available. Since we determined cryo-EM structures of both forms of hTom70, it is worth comparing them with yTom70/71 to highlight notable differences between these homologous proteins. It should be noted that yeast has two Tom70 homologues, yTom70 and yTom71, both referred to as yTom70 hereinafter.

The cryo-EM structure revealed that free hTom70 primarily exists as a monomer, also consistent with SEC-MALS results (**Fig S4B**), while yTom70 showed a dynamic monomer-dimer equilibrium^22^. yTom70 also undergoes open–closed motions of the hydrophobic pocket where Orf9b binds. In contrast, we only observed the open conformation of hTom70. Moreover, the relative orientation of its N- and C-terminal domains differs from that of yTom70 in the open state (**Figs S10B**). These species-specific differences extend to the Hsp90-bound forms as well^23^ (**Fig S10C and S10D**). These differences highlight the importance of considering species-specific structural variations in comparative analyses. Moreover, yTom70 and hTom70 exhibit some functional differences. For example, yeast does not have MAVS, and thus, yTom70 is not involved in MAVS-mediated antiviral responses.

Our integrative approach provides comprehensive estimates for the binary and ternary interactions among Orf9b, Tom70, and Hsp90. Prior to the present study, these measurements had not been available for full-length proteins. While these results explain how Orf9b inhibits the Tom70–Hsp90 interaction, they also underscore the need for further investigation. For example, Hsp90 recruits TBK1 and IRF3 to MAVS via binding to Tom70, which is essential for interferon production upon viral infection^7^. However, there is no structure of the Tom70–Hsp90 complex with TBK1 or IRF3, and it is unknown whether parts of TBK1 or IRF3 interact with the hydrophobic pocket of Tom70. In such a case, the bound Orf9b helix would likely interfere with this process, which we did not examine in this study. Therefore, the overall inhibitory effect of Orf9b may exceed what is inferred solely from reductions in K_D_ or k_on_.

## Materials and Methods

### Protein Expression and Purification

Genes encoding human Hsp90α (residues 1-732), human Tom70 (residues 108-608), and all SARS-CoV-2 Orf9b mutants were codon-optimized for *Escherichia coli* cells and synthesized via Genscript’s gene-synthesis service. All proteins possessed a N-terminal Avi tag, followed by a His_6_ tag, (GGS)_5_ linker, and a TEV cleavage site. For co-expression of Tom70 (residues 108-608) and Orf9b (residues 1-97), the genes were inserted into a pET-Duet vector, where Tom70 was tag-less and Orf9b contained a N-terminal His_6_ tag. Hsp90 was transformed into BL21 Star (DE3) pLysS *E. coli* cells (Invitrogen), whereas all other constructs were transformed into BL21 (DE3) *E. coli* cells (New England Biolabs).

To express Hsp90, cells were grown in medium supplemented with 1% glucose at 37°C and induced with 1 mM IPTG at 15°C for 18 hours after cooling. Frozen cell pellets were resuspended in lysis buffer (20 mM sodium phosphate (pH 8), 500 mM NaCl, 5 mM imidazole, 5 mM DTT, and 2 mM EDTA) and supplemented with protease inhibitor cocktail (ThermoFisher). The cells were then lysed via ultrasonication, clarified by centrifugation, and the supernatant was loaded onto a Ni-INDIGO column (Cube Biotech) pre-equilibrated with lysis buffer. After washing the column with lysis buffer, eluted fractions were collected through a gradient of elution buffer (20 mM NaPi pH 8, 500 mM NaCl, 500 mM Imidazole, 5 mM DTT, and 2 mM EDTA). The fractions containing Hsp90 were dialyzed with TEV protease overnight (20 mM sodium phosphate, pH 8, 100 mM NaCl, 5 mM DTT). After overnight TEV cleavage, the tag was removed after a second round of Ni-INDIGO purification. Hsp90 was then pooled and dialyzed into ion exchange buffer A (20 mM Tris-HCl pH 8, 20 mM NaCl, 2 mM DTT). Hsp90 was then loaded onto a pre-equilibrated HiPrep Q FF 16/10 column (GE Healthcare) and eluted with a salt gradient. Purified Hsp90 was then pooled, concentrated, and buffer exchanged into the needed buffers prior to use or storage. Protein purity was confirmed via SDS-PAGE and all samples were >95% pure. For biotinylated Hsp90, a fraction of Hsp90 was sequestered and did not undergo TEV cleavage. It was then purified via HiPrepQ and then finally biotinylated via the enzymatic protein biotinylation kit (Sigma-Aldrich). Excess biotin was removed via dialysis.

For Tom70 expression, cells were grown to an OD_600_ of 4.0 in Ultra Yield Flasks (Thompson) at 37°C and induced at 1 mM IPTG for 4 hours. Frozen cell pellets were resuspended in lysis buffer (20 mM sodium phosphate pH 8, 500 mM NaCl, 5 mM imidazole, 2 mM DTT) and supplemented with protease inhibitor cocktail (ThermoFisher). The cells were lysed, clarified, and the supernatant was loaded onto a pre-equilibrated His-Trap HP Ni NTA column (GE Healthcare). After washing the column with lysis buffer, eluted fractions were collected through a gradient of elution buffer (20 mM sodium phosphate pH 8, 500 mM NaCl, 500 mM imidazole, 2 mM DTT). Eluted fractions were then dialyzed overnight with TEV protease (20 mM sodium phosphate pH 8, 100 mM NaCl, 5 mM DTT), and the tags were removed by a second Ni column purification. Samples were then further purified by size-exclusion chromatography using a HiLoad 16/600 Superdex 200 pg column (GE Healthcare) with storage buffer (20 M sodium phosphate pH 7, 80 mM NaCl, 2 mM DTT). Protein purity was confirmed via SDS-PAGE and all samples were >95% pure.

For co-expression of Tom70 and Orf9b, cells were grown to an OD_600_ of 4.0 in Ultra Yield Flasks (Thompson) at 37°C and induced at 1 mM IPTG for 4 hours. Frozen cell pellets were resuspended in the lysis buffer and supplemented with protease inhibitor cocktail (ThermoFisher). The cells were lysed, clarified, and the supernatant was loaded onto a pre-equilibrated His-Trap HP Ni NTA column (GE Healthcare). After washing the column with lysis buffer, eluted fractions were collected through a gradient of the elution buffer. Eluted fractions were then purified by size-exclusion chromatography using a HiLoad 16/600 Superdex 200 pg column (GE Healthcare) with the storage buffer. Protein purity was confirmed via SDS-PAGE and all samples were >95% pure.

For expression of all Orf9b mutants, cells were grown to an OD_600_ of 4.0 in Ultra Yield Flasks (Thompson) at 37°C and induced at 1 mM IPTG for 4 hours. Frozen cell pellets were first resuspended in lysis buffer and supplemented with protease inhibitor cocktail (ThermoFisher). The cells were lysed, clarified, and the supernatant was separated from the inclusion body. The inclusion body was then resuspended in denaturing lysis buffer (lysis buffer + 4 M Guanidinium chloride) and loaded onto a pre-equilibrated His-Trap HP Ni NTA column (GE Healthcare). Eluted fractions were then collected through a gradient of the denaturing elution buffer (elution buffer + 4 M Guanidinium Chloride), and extensively dialyzed to remove the denaturant. Orf9b mutants were then subjected to TEV protease, and tags were removed through a second, native His-Trap HP Ni NTA column (GE Healthcare). Samples were then dualized into storage buffer (20 M sodium phosphate pH 7, 80 mM NaCl, 2 mM DTT) and stored. Protein purity was confirmed via SDS-PAGE and all samples were >95% pure.

Hsp90_EEVD_ peptide (Y^724^<u>DTSRMEEVD</u>^732^) and Orf9b_Helix_ peptide (^44^<u>IILRLGSPLSLNMARKTLNSLEDKAFQLTPIAVQ</u>^77^ Y) were ordered from LifeTein. The tyrosine residues were added to aid with quantification. The C-terminus of the Hsp90_EEVD_ peptide was not amidated in order to preserve function of the C-terminus carboxyl group.

### Size Exclusion Chromatography with Multi-Angle Light Scattering (SEC-MALS)

25 µM Tom70, Tom70:Orf9b_Helix_ complex (1:2 ratio), or Tom70:Orf9b complex were incubated for 2 hours at 25°C. The individual proteins or protein complexes were then injected into a Superdex 200 Increase 10/300 GL column (GE Healthcare) connected to a miniDAWN MALS and an Optilab dRI detector (Wyatt Technology), using a running buffer of 25 mM Tris-HCl pH 8, 200 mM NaCl, and 2 mM TCEP. The molecular weight was determined through the ASTRA software package v8.2.2 (Wyatt Technology).

### Isothermal Titration Calorimetry

All ITC experiments were recorded at 20°C using a Microcal-PEAQ-ITC calorimeter (Malvern Panalytical). Experiments involving full-length Hsp90 or the Hsp90_EEVD_ peptide were carried out in buffer containing 20 mM sodium phosphate (pH 8), 150 mM NaCl, 2 mM TCEP, and 10% glycerol. 10% glycerol was required to prevent precipitation of Hsp90 during titration.

For full-length Hsp90 experiments, Tom70 (400 µM), Tom70:Orf9b_Helix_ complex (400 µM complex), and Tom70:Orf9b complex (500 µM) were titrated into Hsp90 (20 µM). For the Hsp90_EEVD_ peptide experiments, the Hsp90_EEVD_ (400-500 µM) was titrated into Tom70, Tom70:Orf9b_Helix_ complex, and Tom70:Orf9b complex (30 µM) in the cell. The Tom70:Orf9b_Helix_ complex was created by mixing Tom70:Orf9b_Helix_ in a 1:2 ratio, and the Tom70:Orf9b complex was produced through coexpression. Unbound Orf9b_Helix_ was removed via multiple rounds of concentration and buffer exchange on a concentrator (Vivaspin 20, Satorius). Orf9b truncation mutant complexes were produced in the same way. Experiments involving the binding of Orf9b_Helix_ titrated into Tom70 were carried out in buffer containing 20 mM sodium phosphate (pH 7), 100 mM NaCl, and 2 mM TCEP. Orf9b_Helix_ (120 µM) was titrated into 15 µM Tom70 or Tom70:Hsp90_EEVD_ complex (15 µM Tom70: 400 µM Hsp90_EEVD_).

All data was fit using a 1:1 binding model to obtain the reported thermodynamic parameters using the software provided by the instrument. Reported parameters are the average and standard deviation of individual runs of the experiment, as outlined in supplementary information (**Figs S3 and S9).**

### Cryo-EM data collection

Cryo-EM samples were first dialyzed into cryo-EM buffer (20 mM HEPES (pH 7), 80 mM NaCl, and 3 mM TCEP) and then concentrated into their final concentrations: 40 µM Tom70 for the free structure, and 80 µM Tom70 with 400 µM Hsp90_EEVD_ peptide for the complex. 3 µL of each sample were applied to a glow discharged grid (200 mesh R 1.2/1.3 Cu, Quantifoil) before being blotted for 5-6s under 100% humidity at 4°C and plunged into liquid ethane using a Vitrobot Mark IV (ThermoFisher). CHAPS was added to a final concentration of 0.05% immediately prior to plunge freezing. Each sample was then imaged on a Titan Krios G4 Electron Microscope (ThermoFisher) operating at 300 kV equipped with a Gatan BioContinuum Imaging Filter. Micrographs were recorded through a Gatan K3 direct electron detector in super-resolution mode at a nominal magnification of 105,000x, corresponding to 0.416 Å/pixel. During processing, images were binned by a factor of 2, corresponding to 0.832 Å/pixel.

### Cryo-EM image processing

Data was processed through the CryoSPARC software package^59^ (v3.3.1, Strucutra Biotechnology). For the free Tom70 structure, a total of 5453 micrographs were collected, motion corrected, and patch CTF estimated. First, an initial set of particles was picked using the blob picker and underwent multiple rounds of 2D classification. The best 2D classes were then provided as templates for template picking for an initial number of 2,627,413 particles. Multiple rounds of 2D classification were performed in order to obtain 372,624 “good” particles. Ab-initio reconstruction was then performed to generate three 3D reconstructions of Tom70. The best reconstruction was then selected, and all particles were used for a non-uniform refinement that resulted in a map of Tom70 at 3.57Å. The half maps from this non-uniform refinement were then sharpened using deepEMhancer^60^. An in-depth workflow can be found in supplementary information (**Fig S5**).

For the Tom70 with Hsp90_EEVD_ peptide structure, a total of 5301 micrographs were collected, motion corrected, and patch CTF estimated. An initial set of particles was picked using the blob picker and underwent multiple rounds of 2D classification. The best 2D classes were then provided as templates for template picking for an initial number of 2,516,289 particles. Multiple rounds of 2D classification were performed in order to obtain 353,087 “good” particles. Ab-initio reconstruction was then performed to generate three 3D reconstructions. The best reconstruction was then selected, and all particles were used for a non-uniform refinement that resulted in a map of Tom70:Hsp90_EEVD_ peptide at 3.64Å. The half maps from this non-uniform refinement were then sharpened using deepEMhancer^60^. An in-depth workflow can be found in supplementary information **(Fig S6).**

### Model Building

Initial models for Tom70 and Tom70:Hsp90_EEVD_ complex were generated through AlphaFold^61,62^, and the initial fitting into the map was performed in UCSF ChimeraX^63^. Each respective model was then refined against each respective map through multiple rounds of real-space refinement in PHENIX^64^ and manual adjustments in COOT^65^. Portions that were not present in the density were deleted and the final quality was assessed using MolProbity^66^. All collection, refinement, and validation statistics are given in supplementary information (**Table S1)**.

### Sample Preparation for ^19^F NMR Spectroscopy

Isotopically labeled Tom70 and Co-expressed Tom70:Orf9b for NMR studies were prepared by growing the respective transformed BL21(DE3) *E.coli* cells in M9 media supplemented with 5-fluoro-tryptophan. Proteins were purified as discussed previously, until >95% pure. After purification, the labeled proteins were prepared in a buffer containing 20 mM sodium phosphate (pH 7), 80 mM NaCl, 2 mM TCEP, 1 mM EDTA, and 10% D_2_O. A typical sample for ^19^F NMR contained ∼ 100 µM ^19^F Tom70. Orf9b_Helix_ and Hsp90_EEVD_ peptides were added to final concentrations of 130 µM and 400 µM respectively **(Fig 3D)**. Hsp90 was added to a final concentration of 200 µM **(Fig S11)**.

Labeling of Orf9b (1-93) with a spin-label was achieved by introducing a single cysteine mutation at V92C. The Orf9b V92C mutant was purified (as discussed above) and prior to labeling, was reduced with 10 mM TCEP for one hour at 25 °C. Orf9b V92C sample was then buffer exchanged (20 mM sodium phosphate (pH 8), 80 mM NaCl) using a HiTrap Desalting column (Cytiva) to remove reducing agent. 30-fold molar excess of MTSL (Sigma-Aldrich) was immediately added to the sample and incubated at 25 °C in the dark for two hours. Orf9b V92C and ^19^F Tom70 were then buffer exchanged (20 mM sodium phosphate (pH 8), 80 mM NaCl, 1 mM EDTA, and 10% D_2_O) and mixed in a 1:1.5 ratio of ^19^F Tom70 and Orf9b V92C to form a complex. The reduced sample was generated through the addition of ascorbic acid (2 mM final concentration).

### ^19^F NMR Acquisition and Processing

NMR experiments were conducted on a Bruker Avance III Neo 600 MHz instrument equipped with a Prodigy cryoprobe (Bruker). The temperature was calibrated using methanol-d_4_ set at 25°C. One dimensional ^19^F NMR spectra were collected using a spectral width of 20 ppm and a ^19^F carrier frequency of-124 ppm. The recycle delay was set to 4s in all experiments with a varied number of scans.

Data processing and analyses were performed using MestReNova (v.14.2.0). The ^19^F chemical shifts were referenced to an external standard, α,α,α-trifluorotoluene (Sigma-Aldrich), which resonates at-63.72 ppm relative to CFCl_3_ at 0 ppm. All data was zero-filled twice and Fourier-transformed using either the 50 Hz Gaussian apodization function **(Figs 4D and S11)** or using the 20 Hz Gaussian apodization **(Fig 3D)**. Peak deconvolutions were fitted using the “line fitting” tool in MestReNova by applying a Lorentzian-Gaussian function^67^.

### Biolayer Interferometry

The binding of surface-immobilized Hsp90 to Tom70, and Tom70:Orf9b was measured at 25°C using a Gator Pilot biolayer interferometry instrument (Gator Bio). Biotinylated Hsp90 proteins were immobilized onto streptavidin biosensor probes, and experiments were conducted in buffer containing 20 mM sodium phosphate (pH 7), 150 mM NaCl, 1% BSA, 0.6 M sucrose, and 1 mM TCEP^68^. Association and dissociation phases were measured for 20s and 100s respectively, except for Tom70:Orf9b, where the association was measured at 5s due to heterogenous binding. To combat heterogenous binding, dissociation curves were fit with a two-phase exponential curve^69^. Due to the association rate (k_on_) being too rapid to quantify, k_on_ values were indirectly estimated by combining k_off_ values obtained from BLI with KD values determined by ITC 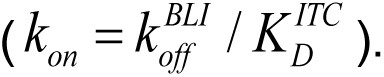 All data are corrected for non-specific binding. Error is given as the standard deviation of at least two measurements.

## Data Availability

All data reported in the main and supplementary data are available upon reasonable request. The electron density maps of Tom70 (EMD-73339) and Tom70 complexed with Hsp90_EEVD_ peptide (EMD-73359) have been deposited in the Electron Microscopy Data Bank (EMDB). The atomic coordinates for Tom70 (PDB: 9YQL) and Tom70:Hsp90_EEVD_ complex (PDB: 9YR4) have been deposited in the Protein Data Bank (PDB).

## Acknowledgements

This work was supported by the NIH grant R35 GM152007 (J.-H.C.), R01 GM108998 (T.I.I.), and the Welch Foundation grant A-2028-20230405 (J.-H.C.). The cryo-EM data was collected at the Laboratory for Biomolecular Structure and Dynamics (LBSD) of Texas A&M University. The LBSD is supported, in part, by the Department of Biochemistry and Biophysics, AgriLife, and Texas A&M University.

