## Supplementary Information for "Molecular mechanism by which SARS-CoV-2 Orf9b suppresses the Tom70-Hsp90 interaction to evade innate immunity"

Figure S1

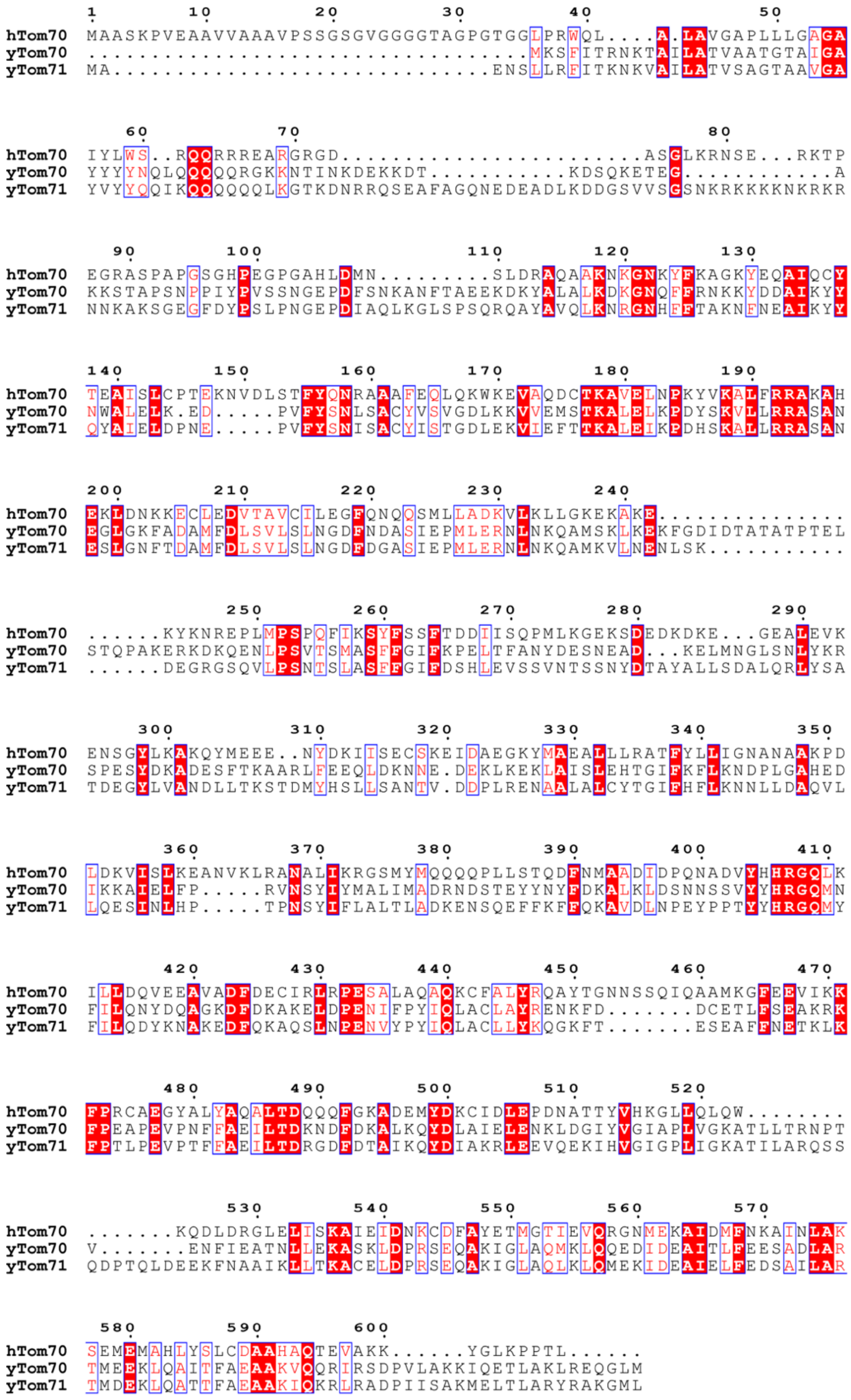


**Supplementary Figure 1:** Alignment of Human Tom70 to yeast Tom70/71. Multiple sequence alignment of Human Tom70 (UniProt ID: O94826), yeast Tom70 (UniProt ID: P07213) , and yeast Tom71 (UniProt ID: P38825) using the Clustal Omega webserver (https://www.ebi.ac.uk/jdispatcher/msa/clustalo) and displayed in Espript 3.0 (https://espript.ibcp.fr/ESPript/ESPript/index.php). Invariant residues are displayed with red background and conservative substitutions are displayed in red font. yTom70 has a 22% sequence identity (69% similarity) and yTom71 has a 21% sequence identify ( 68% similarity) with hTom70. There is an average of 20% identity (68% similarity) between all three sequences.

Figure S2

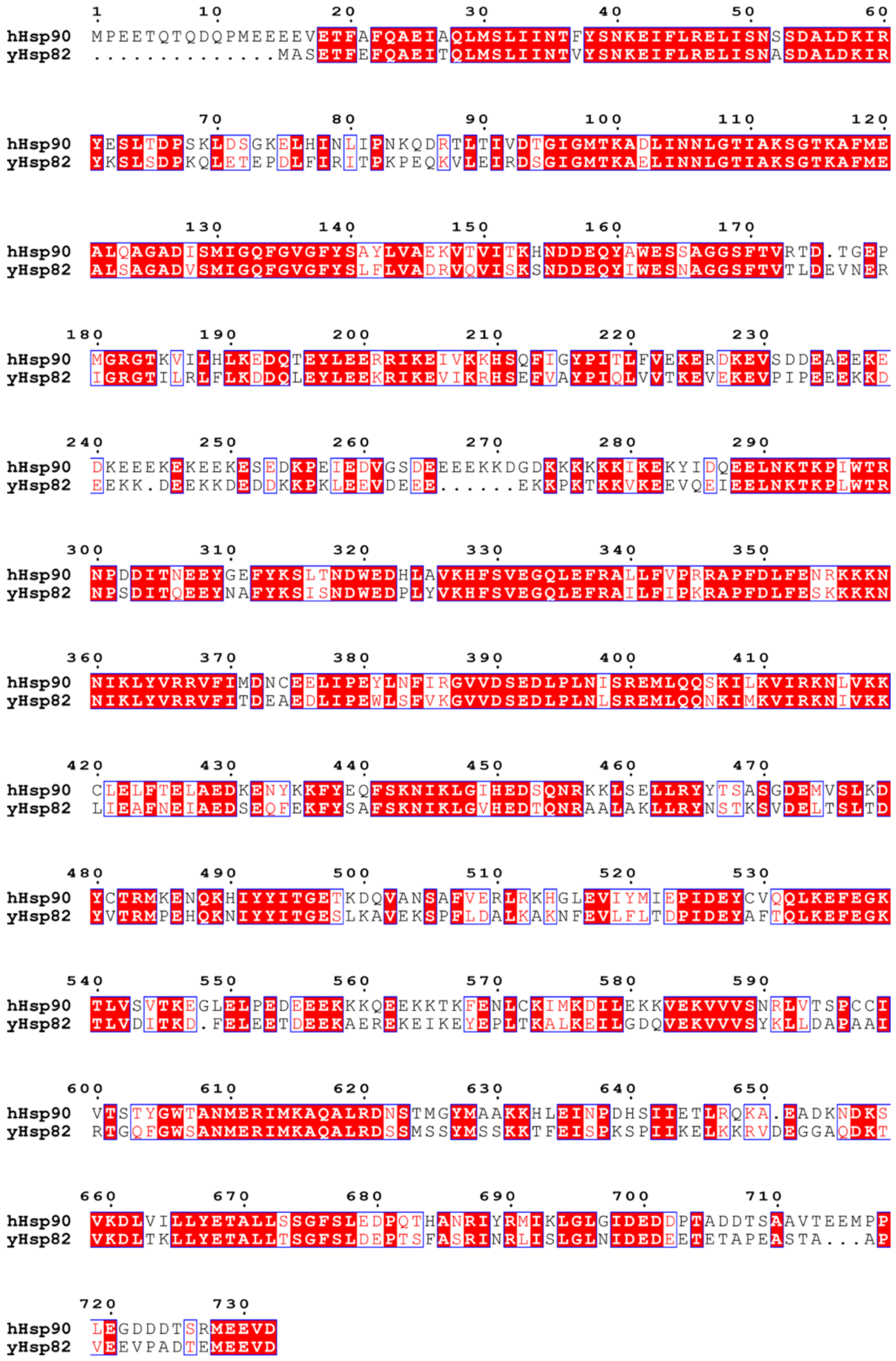


**Supplementary Figure 2:** Alignment of Human Hsp90 to yeast Hsp82. Multiple sequence alignment of Human Hsp90 (UniProt ID: P07900) and yeast Hsp82 (UniProt ID: P02829) using the Clustal Omega webserver (https://www.ebi.ac.uk/jdispatcher/msa/clustalo) and displayed in Espript 3.0 (https://espript.ibcp.fr/ESPript/ESPript/index.php). Invariant residues are displayed with red background and conservative substitutions are displayed in red font. On average, hHsp90 and yHsp90 share 60% sequence identity and 87% sequence similarity.

Figure S3

A

**Hsp90 to Tom70**

Run 1 | Run 2 | Run 3

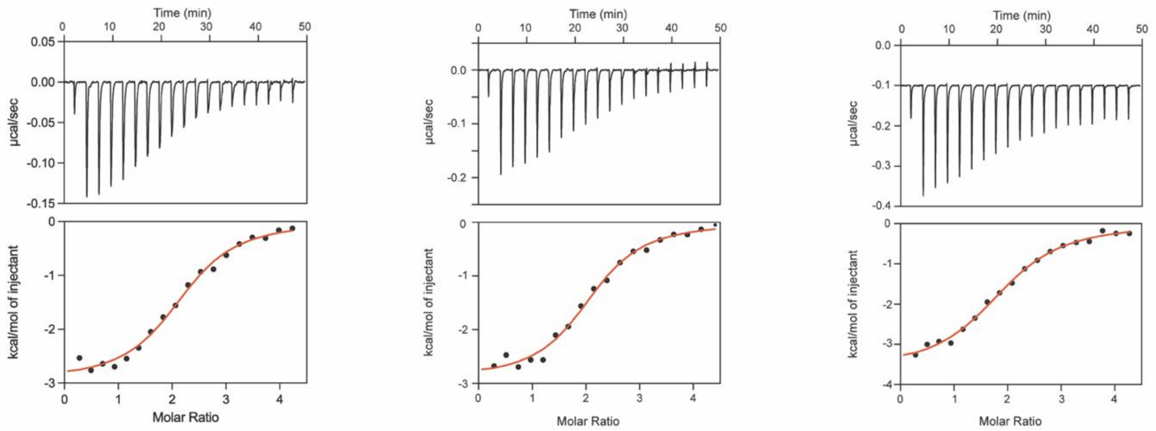


| | Run 1 | Run 2 | Run 3 | Average |
|---|---|---|---|---|
| **Stoichiometry (n)** | 2.18 ± 0.07 | 2.08 ± 0.06 | 1.93 ± 0.05 | 2.06 ± 0.13 |
| **$K_D$ (µM)** | 2.53 ± 0.65 | 2.41 ± 0.63 | 3.96 ± 0.76 | 2.97 ± 0.86 |
| **ΔH (kcal/mol)** | -2.95 ± 0.18 | -2.91 ± 0.16 | -3.62 ± 0.22 | -3.16 ± 0.40 |
| **-TΔS (kcal/mol)** | -4.56 ± 0.22 | -4.62 ± 0.22 | -3.62 ± 0.05 | -4.27 ± 0.56 |
| **ΔG (kcal/mol)** | -7.51 ± 0.26 | -7.56 ± 0.26 | -7.25 ± 0.19 | -7.44 ± 0.17 |

B

**Hsp90 to Tom70:Orf9b**

Run 1 | Run 2 | Run 3

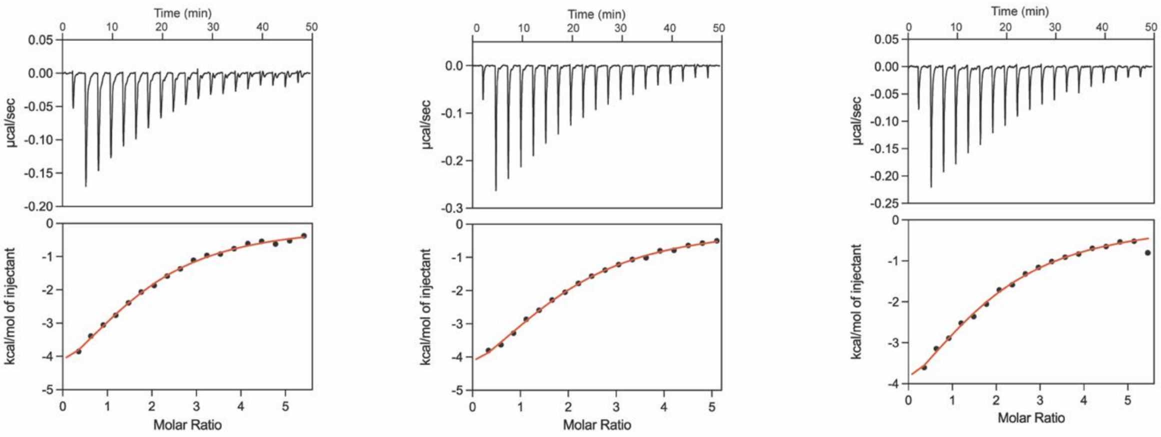


| | Run 1 | Run 2 | Run 3 | Average |
|---|---|---|---|---|
| **Stoichiometry (n)** | 1.71 ± 0.08 | 1.79 ± 0.06 | 1.75 ± 0.09 | 1.75 ± 0.04 |
| **$K_D$ (µM)** | 20.00 ± 4.56 | 25.60 ± 4.58 | 24.00 ± 5.87 | 23.20 ± 2.88 |
| **ΔH (kcal/mol)** | -6.60 ± 0.99 | -7.27 ± 0.81 | -6.61 ± 1.02 | -6.83 ± 0.38 |
| **-TΔS (kcal/mol)** | 0.29 ± 1.00 | 1.11 ± 0.82 | 0.42 ± 1.03 | 0.61 ± 0.44 |
| **ΔG (kcal/mol)** | -6.31 ± 0.23 | -6.16 ± 0.18 | -6.19 ± 0.25 | -6.22 ± 0.08 |

Figure S3 (continued)

C

**Hsp90 to Tom70:Orf9b$_{Helix}$**

Run 1 Run 2 Run 3

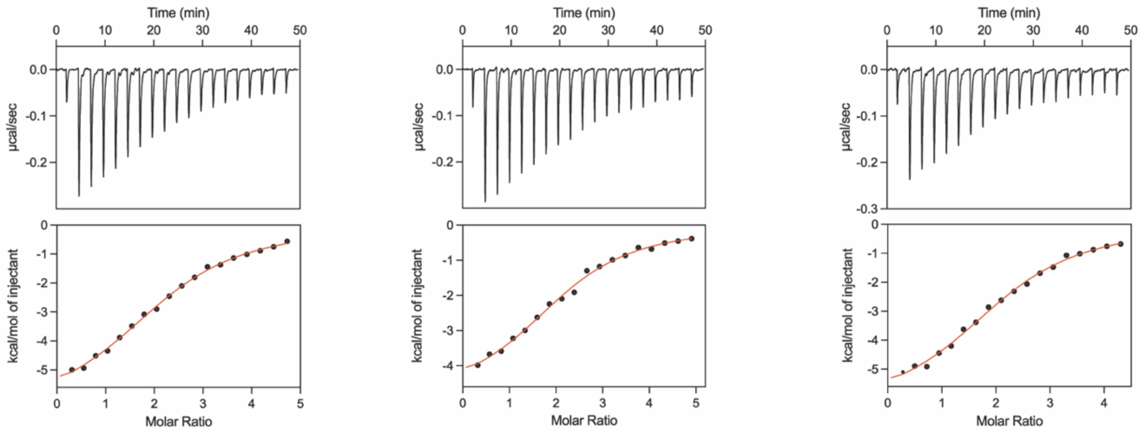


| | Run 1 | Run 2 | Run 3 | Average |
|---|---|---|---|---|
| **Stoichiometry (n)** | 2.24 ± 0.07 | 1.8 ± 0.06 | 2.05 ± 0.08 | 2.03 ± 0.22 |
| **$K_D$ (µM)** | 10.30 ± 1.78 | 7.24 ± 1.67 | 8.00 ± 1.75 | 8.51 ± 1.59 |
| **ΔH (kcal/mol)** | -6.51 ± 0.42 | -5.88 ± 0.48 | -6.70 ± 0.52 | -6.36 ± 0.43 |
| **-TΔS (kcal/mol)** | -0.19 ± 0.44 | -1.02 ± 0.49 | -0.14 ± 0.54 | -0.45 ± 0.56 |
| **ΔG (kcal/mol)** | -6.69 ± 0.17 | -6.90 ± 0.23 | -6.84 ± 0.22 | -6.36 ± 0.43 |

D

**Hsp90 to Tom70:Orf9b ΔCDT (1-80)**

Run 1 Run 2

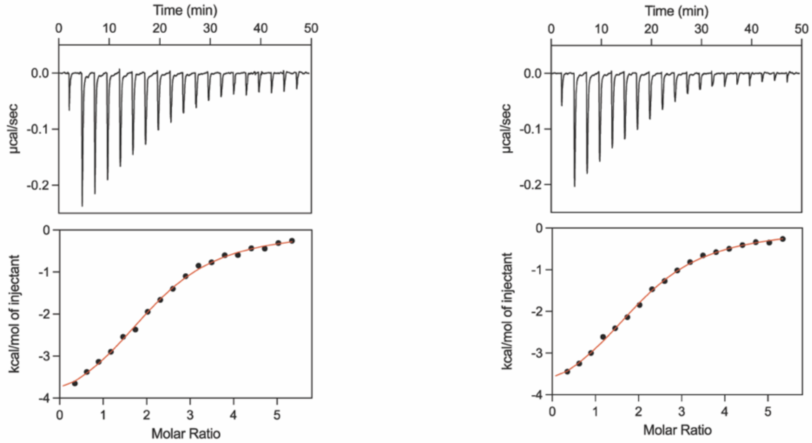


| | Run 1 | Run 2 | Average |
|---|---|---|---|
| **Stoichiometry (n)** | 2.13 ± 0.04 | 2.05 ± 0.04 | 2.09 ± 0.06 |
| **$K_D$ (µM)** | 8.77 ± 1.23 | 9.27 ± 1.12 | 9.02 ± 0.35 |
| **ΔH (kcal/mol)** | -4.52 ± 0.22 | -4.4 ± 0.19 | -4.46 ± 0.08 |
| **-TΔS (kcal/mol)** | -2.27 ± 0.23 | -2.35 ± 0.20 | -2.31 ± 0.06 |
| **ΔG (kcal/mol)** | -6.78 ± 0.14 | -6.75 ± 0.12 | -6.77 ± 0.02 |

Figure S3 (continued)

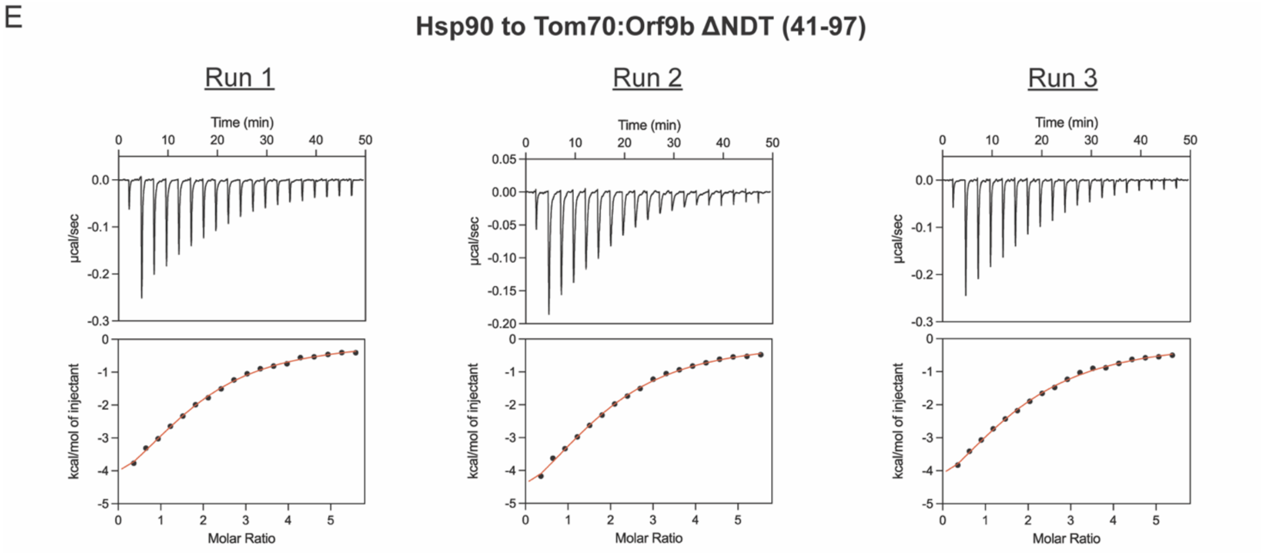


| | Run 1 | Run 2 | Run 3 | Average |
|---|---|---|---|---|
| Stoichiometry (n) | 1.76 ± 0.05 | 1.81 ± 0.06 | 1.75 ± 0.06 | 1.77 ± 0.03 |
| $K_D$ (µM) | 18.10 ± 2.71 | 19.10 ± 3.17 | 22.50 ± 4.15 | 19.90 ± 2.31 |
| ΔH (kcal/mol) | -6.16 ± 0.51 | -6.81 ± 0.62 | -6.82 ± 0.77 | -6.60 ± 0.38 |
| -TΔS (kcal/mol) | -0.29 ± 0.52 | 0.48 ± 0.63 | 0.58 ± 0.78 | 0.26 ± 0.48 |
| ΔG (kcal/mol) | -6.36 ± 0.15 | -6.33 ± 0.17 | -6.23 ± 0.18 | -6.31 ± 0.07 |

**Supplementary Figure 3:** ITC thermograms, isotherms, and thermodynamic parameters for the interaction of full-length Hsp90 with **(A)** free Tom70, **(B)** Tom70:Orf9b complex, **(C)** Tom70:Orf9b$_{Helix}$ complex, **(D)** Tom70:Orf9b ΔCDT (1-80) complex, and **(E)** Tom70:Orf9b ΔNDT (41-97) complex. Numbers after ± symbol under each individual run represent fitting error. Numbers after ± symbol in the average values represent the standard deviation of the repeated runs.

Figure S4

A

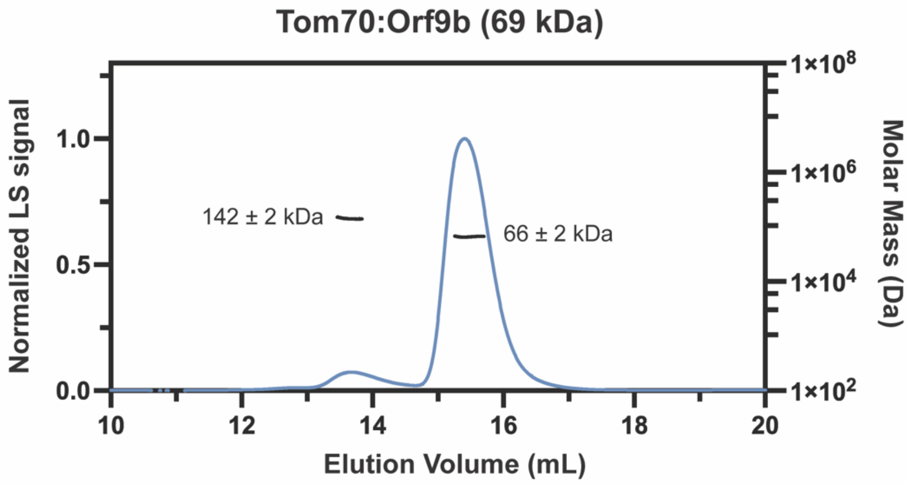


B

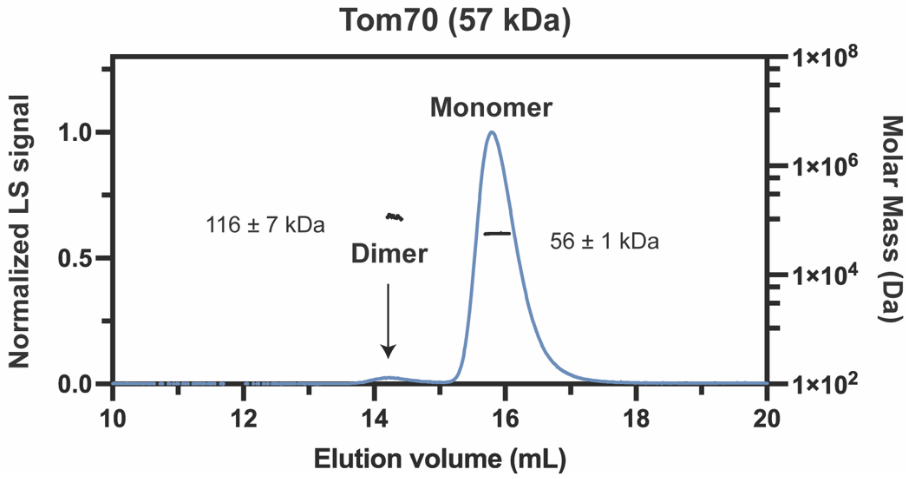


**Supplementary Figure 4:** SEC-MALS chromatogram of **(A)** Tom70-Orf9b complex and **(B)** free Tom70. The theoretical molecular weight of a monomer of Tom70 is 57 kDa and a dimer of Tom70 is 114 kDa. The theoretical molecular weight of Orf9b is 12 kDa making a 1:1 complex 69 kDa.

Figure S5

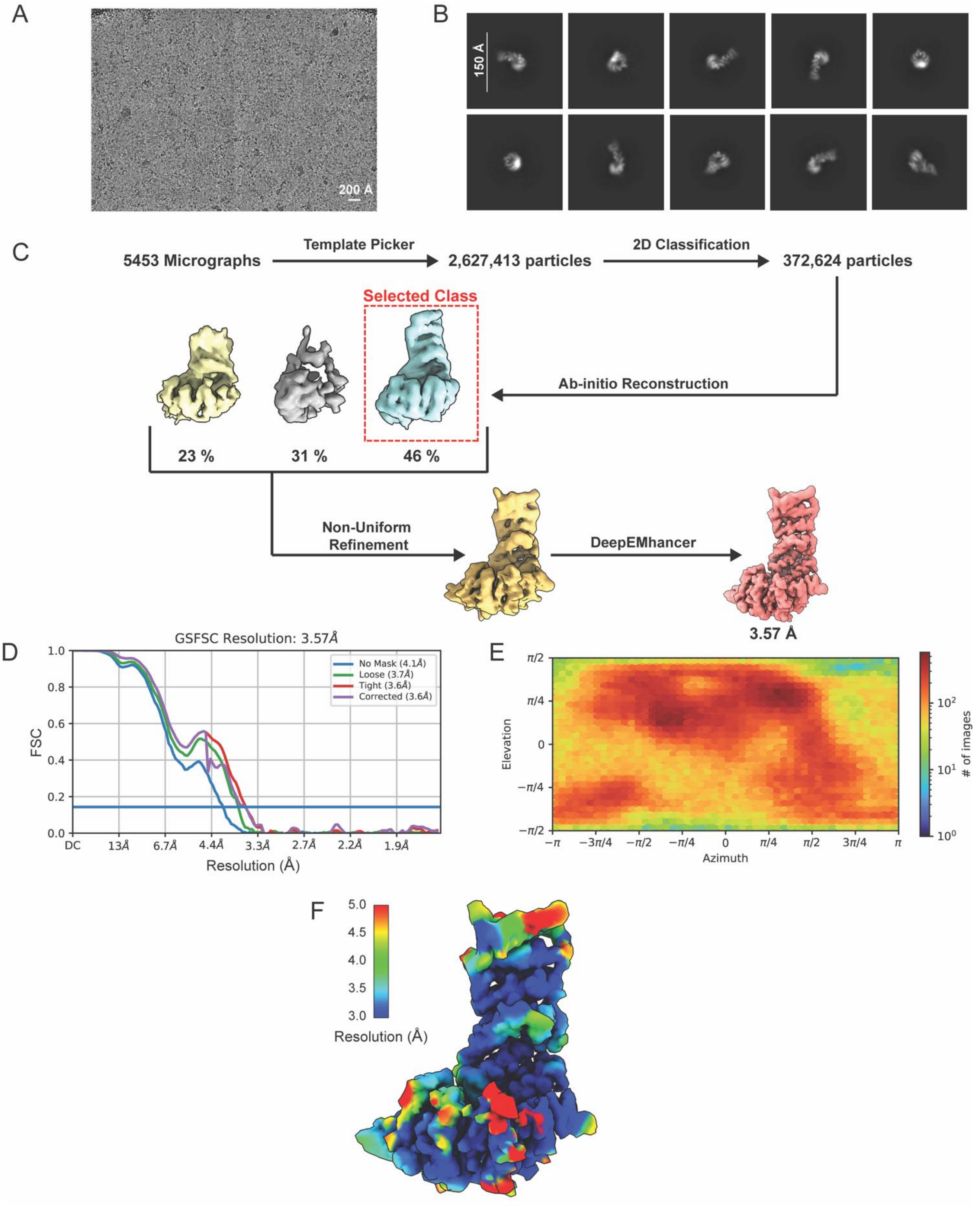


**Supplementary Figure 5: (A)** Representative micrograph, **(B)** representative 2D class averages, **(C)** data processing flow chart, **(D)** Fourier shell correlation (FSC) curves, **(E)** angular distribution, and **(F)** local resolution colored map of the cryo-EM derived Tom70 map.

Figure S6

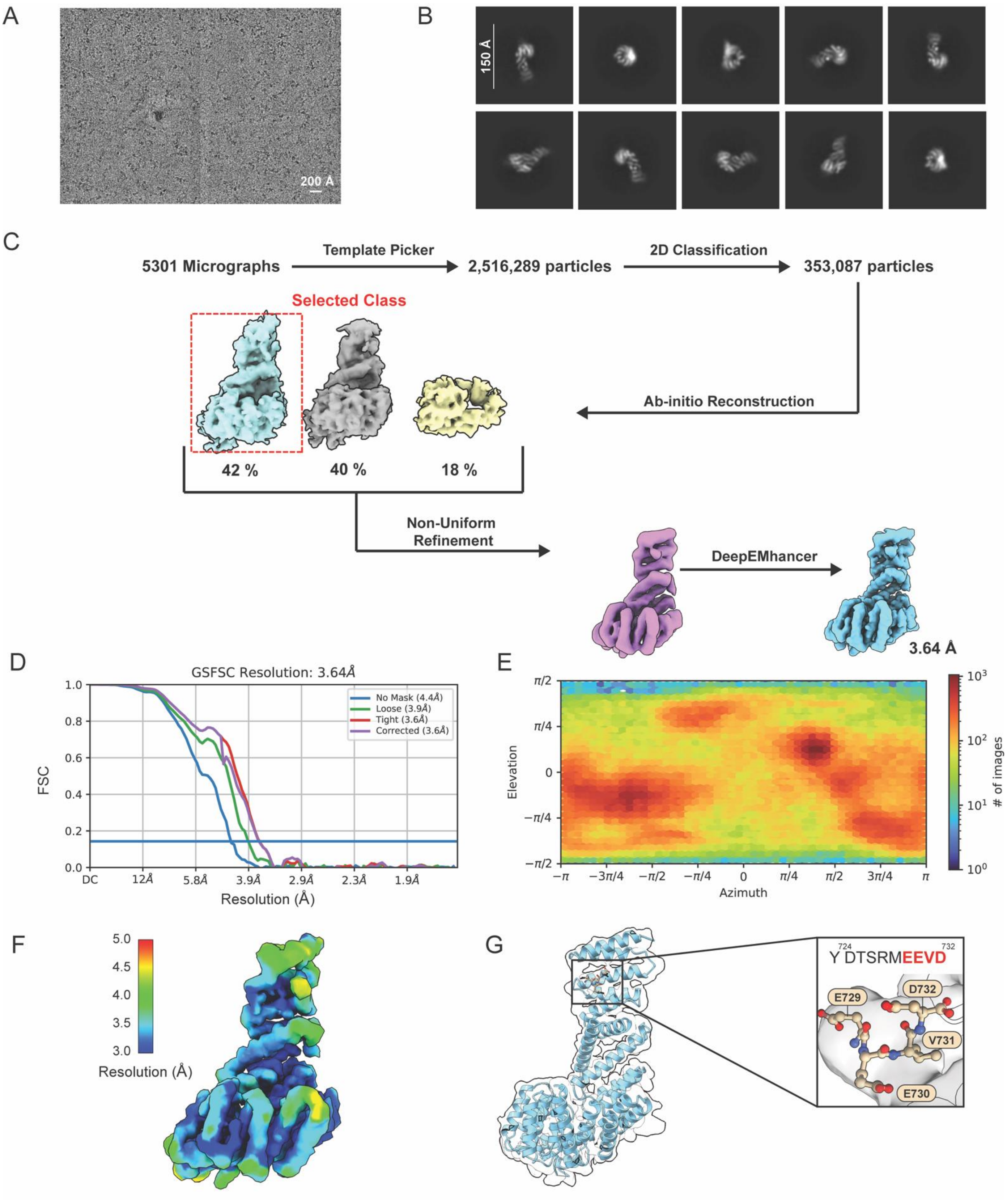


**Supplementary Figure 6: (A)** Representative micrograph, **(B)** representative 2D class averages, **(C)** data processing flow chart, **(D)** Fourier shell correlation (FSC) curves, **(E)** angular distribution, and **(F)** local resolution colored map of the cryo-EM derived Tom70:Hsp90$_{EEVD}$ complex map. **(G)** Only 4 of the 10 residues of the Hsp90$_{EEVD}$ peptide are visualized in the cryo-EM density.

Figure S7

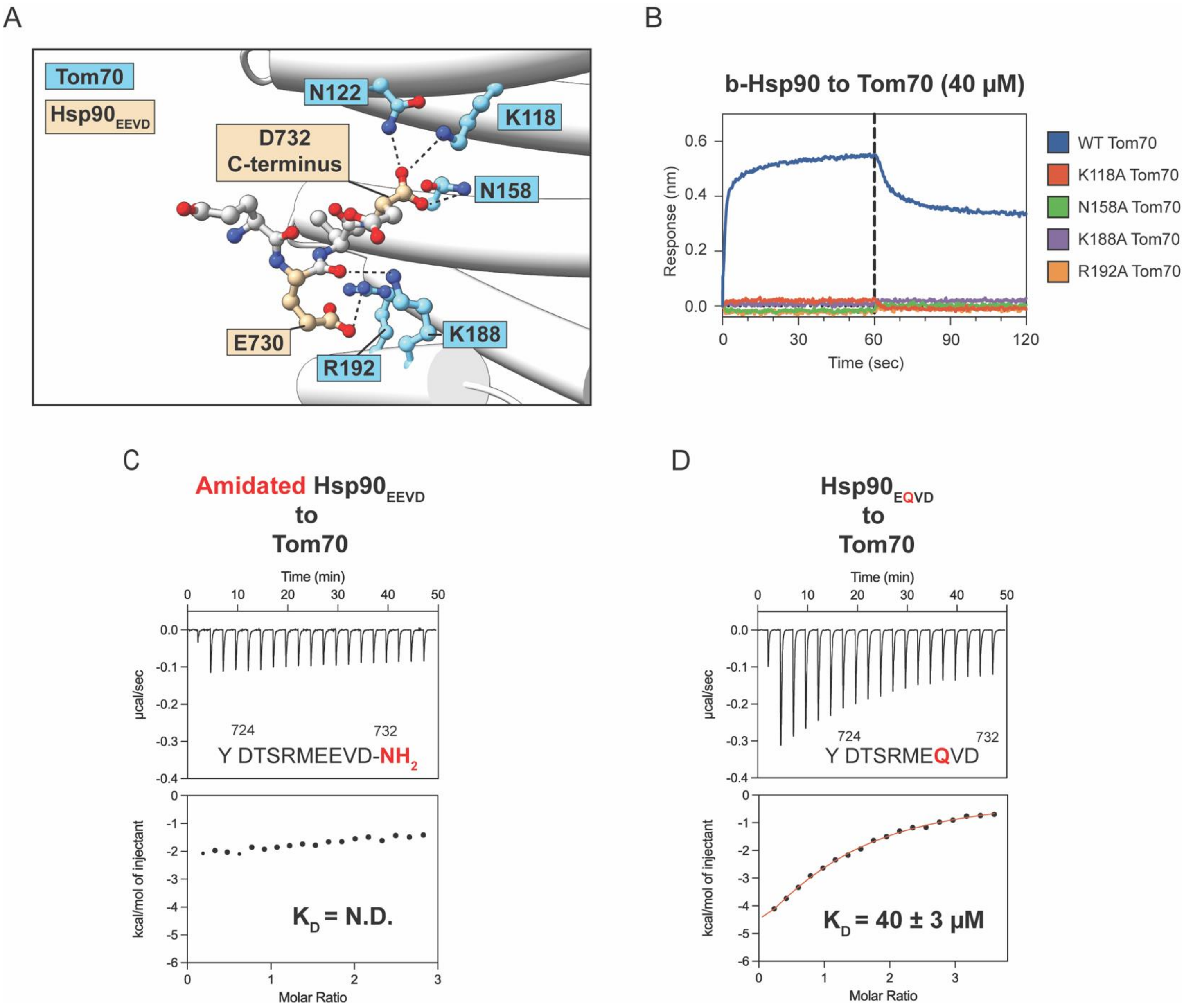


**Supplementary Figure 7: (A)** Interacting residues between Tom70 and $Hsp90_{EEVD}$ peptide. **(B)** BLI sensorgrams of biotinylated full-length Hsp90 assayed to Tom70 mutants. WT refers to "wild-type" Tom70. All Tom70 concentrations are at 40 µM. Dotted line demarcates the beginning of the dissociation phase. **(C-D)** ITC thermograms and isotherms for the interaction of Tom70 to an **(C)** amidated $Hsp90_{EEVD}$ peptide and a **(D)** $Hsp90_{EQVD}$ peptide. N.D. stands for "not-detected" due to the lack of sufficient heat generated.

Figure S8

A

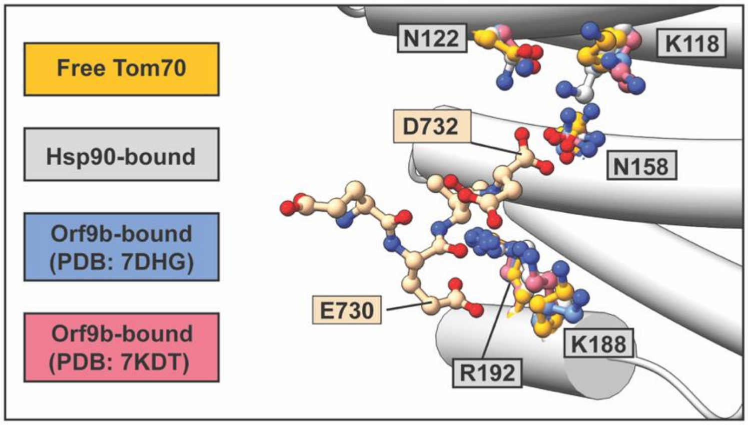


B

| | Hsp90 bound | | Free Tom70 | | Orf9b-bound (PDB: 7DHG) | | Orf9b-bound (PDB: 7KDT) | |
|---|---|---|---|---|---|---|---|---|
| Residue | Backbone SASA (Å²) | Side Chain SASA (Å²) | Backbone SASA (Å²) | Side Chain SASA (Å²) | Backbone SASA (Å²) | Side Chain SASA (Å²) | Backbone SASA (Å²) | Side Chain SASA (Å²) |
| K118 | 0.89 | 45.67 | 0.22 | 52.39 | 0 | 33.80 | 0 | 55.85 |
| N122 | 0 | 26.05 | 3.39 | 41.87 | 0.24 | 42.03 | 0.51 | 42.4 |
| N158 | 4.35 | 27.92 | 3.68 | 24.73 | 1.29 | 26.58 | 0.79 | 34.67 |
| K188 | 4.24 | 105.16 | 7.05 | 119.46 | 4.39 | 122.33 | 7.05 | 126.02 |
| R192 | 3.10 | 43.06 | 2.25 | 47.17 | 0.18 | 47.42 | 0.19 | 43.51 |

C

| Interaction Sites | | Distance (Å) | | | |
|---|---|---|---|---|---|
| Tom70 residue | Hsp90 residue | Hsp90 bound | Free Tom70 | Orf9b-bound (PDB: 7DHG) | Orf9b-bound (PDB: 7KDT) |
| K 118 NZ | D 732 OXT | 3.26 | 7.33 | 6.14 | 6.72 |
| N 122 ND2 | D 732 OX | 2.87 | 3.44 | 3.45 | 3.45 |
| N 158 ND2 | D 732 O | 3.88 | 4.22 | 3.60 | 3.18 |
| K 188 NZ | E 730 O | 3.22 | 5.19 | 6.56 | 3.67 |
| K 188 NZ | E 730 OE2 | 3.92 | 5.27 | 5.89 | 4.52 |
| R 192 NH2 | E 730 OE1 | 4.11 | 4.35 | 4.01 | 4.04 |

**Supplementary Figure 8: (A)** The side chains involved in $Hsp90_{EEVD}$ binding for free Tom70, $Hsp90_{EEVD}$ bound Tom70, and the two available structures of Orf9b-bound Tom70 are shown. The N-terminal region of each Tom70 was aligned (through matchmaker in ChimeraX) to the $Hsp90_{EEVD}$-bound form in order to calculate pairwise distance. **(B)** Calculated Solvent Assessable Surface Area (SASA) for each bound state of Tom70. SASA was calculated through the GETAREA software (https://curie.utmb.edu/getarea.html#:~:text=Sealy%20Center%20for%20Structural%20Biology,.edu%20or%20webraunATutmb.edu%20.). **(C)** Pairwise distance measurements for the interacting residues of Tom70 and Hsp90 in each bound state of Tom70.

Figure S9

A

**$Hsp90_{EEVD}$ to Tom70**

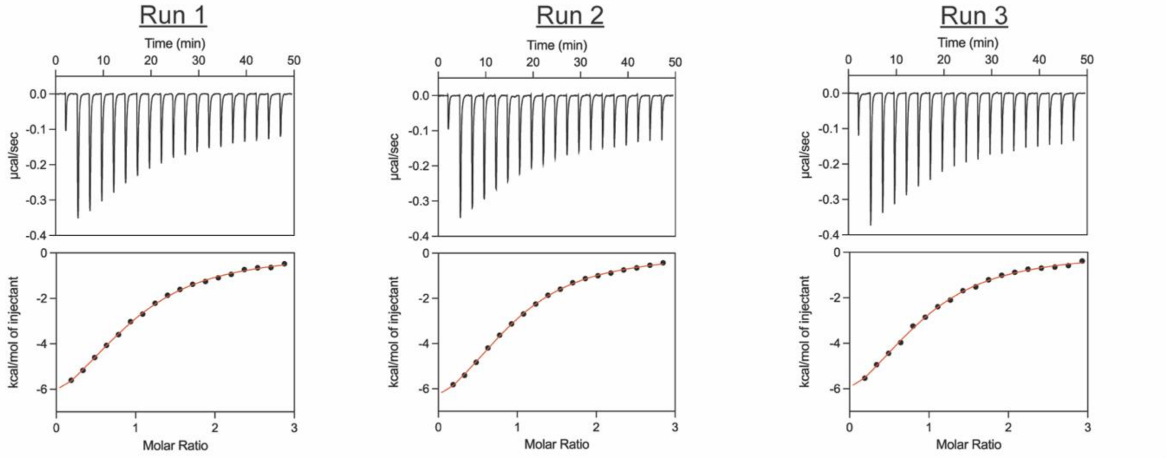


| | Run 1 | Run 2 | Run 3 | Average |
|---|---|---|---|---|
| **Stoichiometry (n)** | 0.94 ± 0.02 | 0.92 ± 0.01 | 0.89 ± 0.02 | 0.91 ± 0.03 |
| **$K_D$ (µM)** | 12.80 ± 1.43 | 10.90 ± 0.83 | 11.90 ± 1.65 | 11.87 ± 0.95 |
| **ΔH (kcal/mol)** | -8.82 ± 0.50 | -8.79 ± 0.32 | -8.75 ± 0.63 | -8.79 ± 0.04 |
| **-TΔS (kcal/mol)** | 2.26 ± 0.51 | 2.13 ± 0.32 | 2.14 ± 0.63 | 2.18 ± 0.07 |
| **ΔG (kcal/mol)** | -6.57 ± 0.11 | -6.66 ± 0.08 | -6.61 ± 0.14 | -6.61 ± 0.05 |

B

**$Hsp90_{EEVD}$ to $Tom70:Orf9b_{Helix}$**

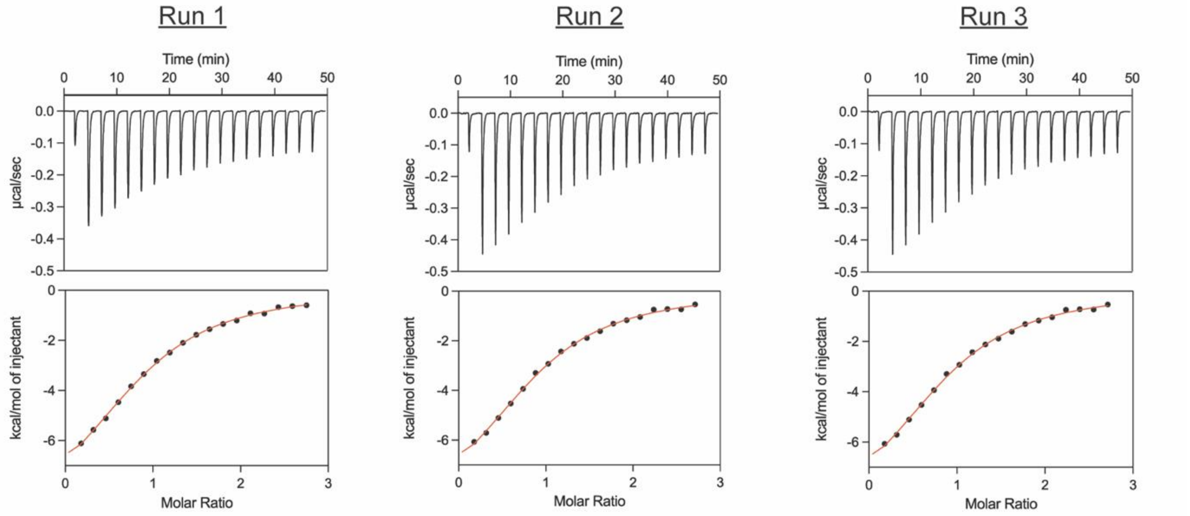


| | Run 1 | Run 2 | Run 3 | Average |
|---|---|---|---|---|
| **Stoichiometry (n)** | 0.90 ± 0.02 | 0.90 ± 0.02 | 0.93 ± 0.03 | 0.91 ± 0.02 |
| **$K_D$ (µM)** | 12.40 ± 1.46 | 11.80 ± 1.79 | 12.80 ± 1.68 | 12.33 ± 0.50 |
| **ΔH (kcal/mol)** | -9.65 ± 0.58 | -9.49 ± 0.71 | -9.65 ± 0.73 | -9.60 ± 0.09 |
| **-TΔS (kcal/mol)** | 3.07 ± 0.59 | 2.88 ± 0.72 | 3.09 ± 0.73 | 3.01 ± 0.12 |
| **ΔG (kcal/mol)** | -6.58 ± 0.12 | -6.61 ± 0.16 | -6.57 ± 0.13 | -6.59 ± 0.02 |

Figure S9 (continued)

C

**Orf9b$_{Helix}$ to Tom70**

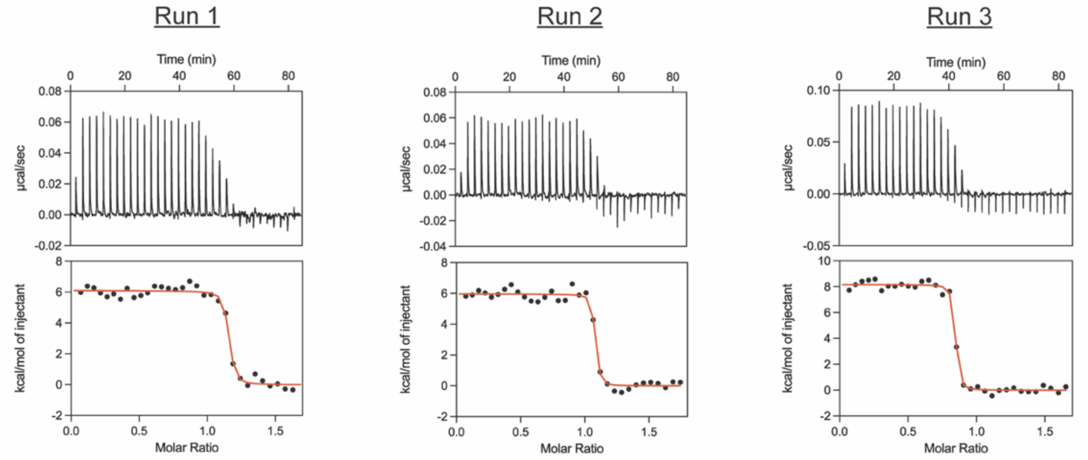


| | Run 1 | Run 2 | Run 3 | Average |
|---|---|---|---|---|
| **Stoichiometry (n)** | 1.13 ± 0.01 | 1.06 ± 0.01 | 0.82 ± 0.01 | 1.00 ± 0.16 |
| **$K_D$ (nM)** | 4.24 ± 1.88 | 2.03 ± 1.21 | 2.14 ± 1.06 | 2.80 ± 1.24 |
| **ΔH (kcal/mol)** | 6.10 ± 0.15 | 5.96 ± 0.13 | 8.16 ± 0.11 | 6.74 ± 1.23 |
| **-TΔS (kcal/mol)** | -17.30 ± 0.30 | -17.60 ± 0.37 | -19.80 ± 0.31 | -18.23 ± 1.37 |
| **ΔG (kcal/mol)** | -11.20 ± 0.44 | -11.70 ± 0.60 | -11.60 ± 0.50 | -11.50 ± 0.26 |

D

**Orf9b$_{Helix}$ to Tom70:Hsp90$_{EEVD}$**

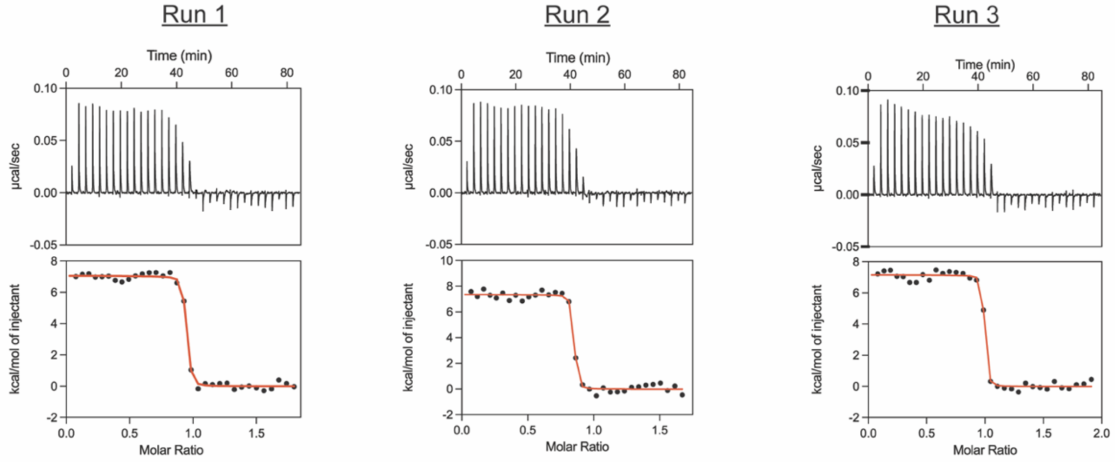


| | Run 1 | Run 2 | Run 3 | Average |
|---|---|---|---|---|
| **Stoichiometry (n)** | 0.92 ± 0.01 | 0.83 ± 0.01 | 0.97 ± 0.01 | 0.91 ± 0.07 |
| **$K_D$ (nM)** | 2.40 ± 0.07 | 1.44 ± 1.03 | 1.05 ± 0.08 | 1.63 ± 0.70 |
| **ΔH (kcal/mol)** | 7.05 ± 0.08 | 7.34 ± 0.11 | 7.15 ± 0.09 | 7.18 ± 0.14 |
| **-TΔS (kcal/mol)** | -18.60 ± 0.08 | -19.20 ± 0.43 | -19.20 ± 0.10 | -19.00 ± 0.35 |
| **ΔG (kcal/mol)** | -11.60 ± 0.03 | -11.90 ± 0.72 | -12.00 ± 0.08 | -11.83 ± 0.21 |

**Supplementary Figure 9:** ITC thermograms, isotherms, and thermodynamic parameters for the interaction of Hsp90$_{EEVD}$ with **(A)** free Tom70 and **(B)** Tom70:Orf9b$_{Helix}$ complex, as well as Orf9b$_{Helix}$ with **(C)** free Tom70 and **(D)** Tom70:Hsp90$_{EEVD}$ complex. Numbers after ± symbol under each individual run represent fitting error. Numbers after ± symbol in the average values represent the standard deviation of the repeated runs.

Figure S10

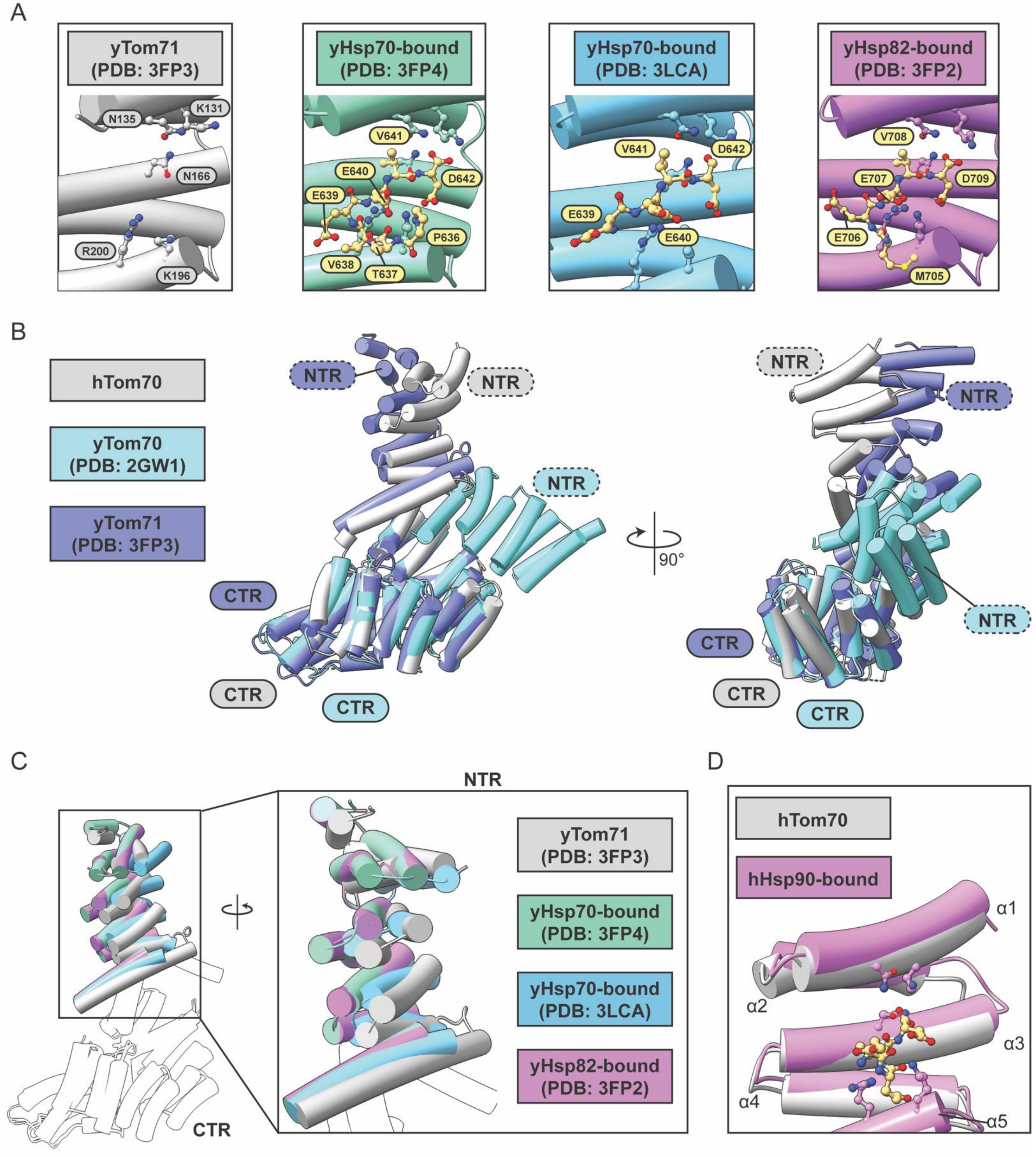


**Supplementary Figure 10: (A)** Comparison of the yHsp70/82 binding site in free yTom71 and all Hsp70/82-bound forms. **(B)** Structural alignment of free human (hTom70) and the two yeast Tom70 homologues (yTom70/71). The N-terminal Domain (NTD) and C-terminal Domain (CTD) are labeled for each. **(C)** All available structures of yTom71 bound yHsp70/82s aligned to free yTom71. The inset displays the NTD. **(D)** hHsp90-bound hTom70 aligned to free hTom70. Only helices α1-α5 are shown.

Figure S11

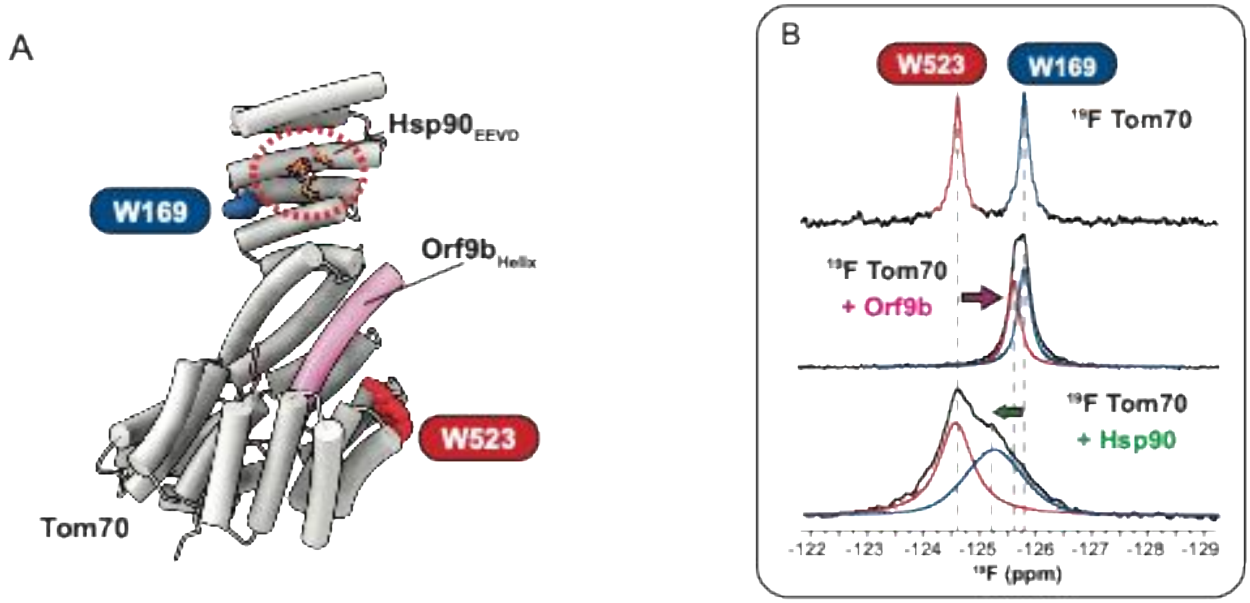


**Supplementary Figure 11: (A)** Schematic for $^{19}$F Tom70. W169 (blue) is proximal to the $Hsp90_{EEVD}$ binding site, and W523 (red) is proximal to the $Orf9b_{Helix}$ binding site on $^{19}$F Tom70. **(B)** $^{19}$F Tom70 spectra for the binding full-length Orf9b (middle) and full-length Hsp90 (bottom) to Tom70.

Figure S12

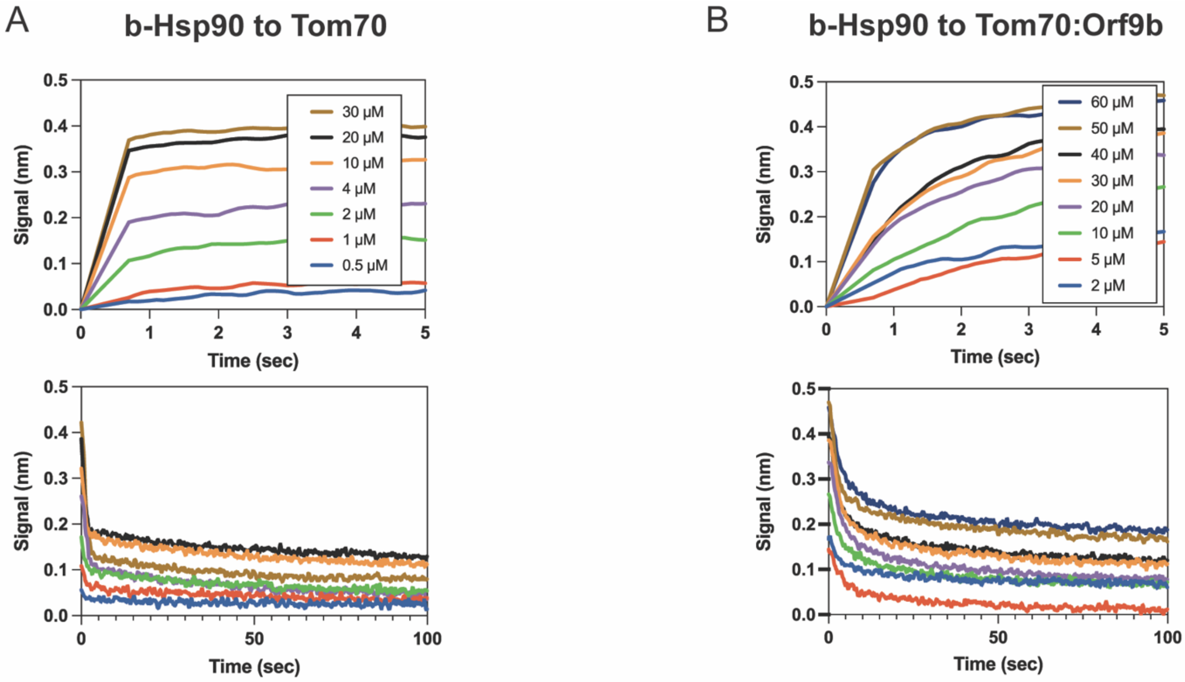


C

| Ligand | Analytes | $k_{off}^{BLI}$ ($s^{-1}$) | $K_D^{ITC}$ (μM) | $k_{on}$ ($\mu M^{-1}s^{-1}$) |
|---|---|---|---|---|
| b-Hsp90 | Tom70 | 1.39 ± 0.38 | 3 ± 1 | 0.46 ± 0.20 |
| | Tom70:Orf9b | 0.39 ± 0.07 | 22 ± 1 | 0.02 ± 0.01 |

**Supplementary Figure 12: (A-B)** Representative BLI sensorgrams of the association (top) and dissociation (bottom) of biotinylated Hsp90 (b-Hsp90) to **(A)** Tom70 and (**B)** Tom70 complexed with full-length Orf9b. (**C**) Table for BLI derived $k_{off}$ values and calculated $k_{on}$ values. $k_{on}$ values were estimated by combining $k_{off}$ values obtained from BLI with $K_D$ values determined by ITC, i.e., $k_{on} = k_{off}^{BLI} / K_D^{ITC}$ . Numbers after the ± represent propagated standard deviation.

Figure S13

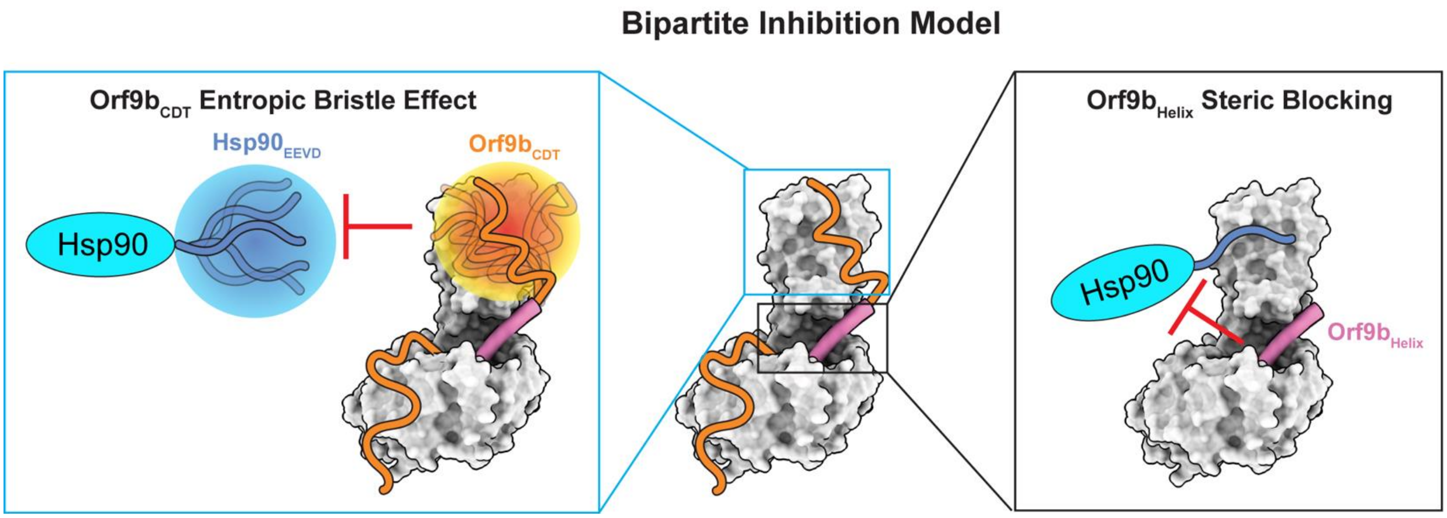


**Supplementary Figure 13:** Orf9b inhibits Hsp90 binding to Tom70 through a dual mechanism. The Orf9b C-terminal disordered tail (CDT) acts as an entropic bristle to sterically hinder the binding of $Hsp90_{EEVD}$ to Tom70. In addition, the $Orf9b_{Helix}$ sterically blocks the ancillary Hsp90 binding site. Orf9b-bound Tom70 (PDB 7DHG) is used to model the bipartite inhibition model.

Table S1

| | Tom70 (EMD-73339) (PDB 9YQL) | $Hsp90_{EEVD}$-bound Tom70 (EMD-73359) (PDB 9YR4) |
|---|---|---|
| **Data collecting and processing** | | |
| Magnification | 105,000 | |
| Voltage (kV) | 300 | |
| Electron exposure ($e^-/Å^2$) | 50 | |
| Defocus range (μM) | -0.8 to -2.4 | |
| Pixel size (Å) | 0.832 | |
| Symmetry Parameters | C1 | |
| Micrographs used | 5453 | 5301 |
| Initial Particle images (no.) | 2627413 | 2516289 |
| Final Particle images (no.) | 372624 | 353087 |
| Map resolution (Å) at FSC threshold 0.143 | 3.57 | 3.64 |
| Map resolution range (Å) | 3.0 to 6.5 | 2.5 to 5.5 |
| Map post-processing | DeepEmhancer | |
| **Refinement** | | |
| Initial model used | AlphaFold2 | |
| Composition | | |
| Non-hydrogen atoms | 3794 | 3819 |
| Protein Residues | 476 | 479 |
| B factors ($Å^2$) | | |
| Protein | 69.36/138.93/94.06 | 81.69/237.99/144.34 |
| R.M.S.D from ideal values | | |
| Length (Å) | 0.004 | 0.004 |
| Bond angles (°) | 0.706 | 0.778 |
| MolProbity score | 1.02 | 1.03 |
| Clashscore | 2.38 | 2.5 |
| Ramachandran plot (%) | | |
| Favored | 98.3 | 98.3 |
| Allowed | 1.7 | 1.7 |
| Disallowed | 0 | 0 |
| CaBLAM outliers (%) | 0.6 | 0.4 |
| Rotamer outliers (%) | 0.3 | 0.3 |
| C-beta outliers (%) | 0.0 | 0.0 |

**Supplementary Table 1:** Cryo-EM data collection, refinement, and validation statistics for Tom70 and $Hsp90_{EEVD}$-bound Tom70 maps and models.
